# Neural cell state modulation by *PARK2* and dopaminergic neuroprotection by small molecule Parkin agonism

**DOI:** 10.64898/2026.04.01.715918

**Authors:** Yongxing Gong, Armin Bayati, Tyler J. Alban, Prerana Parthasarathy, Fang Zhou, Vladimir Makarov, Yuhang Zhao, Chienwen Su, Jackson G. Schumacher, Vijay Singh, Laura A. Volpicelli-Daley, Wen Luo, Thomas M. Durcan, Shovan Dutta, Michael A. Schwarzschild, Jennifer A. Johnston, Timothy A. Chan, Xiqun Chen

## Abstract

Parkin, an E3 ubiquitin ligase encoded by *PARK2*, plays a key role in both hereditary and sporadic Parkinson’s disease (PD), yet there are no therapies currently available that can target this important pathway. Here, we show that Parkin is critical for successful neuronal differentiation and survival, and we develop small-molecule Parkin agonists that can protect dopaminergic neurons. Upon differentiation of neural progenitor cells, loss of Parkin results in a reduced capacity to maintain neuronal cell state, dopaminergic neuronal phenotypes, and stress resistance. Moreover, Parkin loss disrupted cell morphology and the stability of neurites. Transcriptional and single-cell analyses reveal that Parkin controls critical pathways regulating stem-like cell transitions and is needed for stable neuronal maturation. We also examined the effects of FB231, a small molecule enhancer of Parkin E3 ligase activity, in models of PD. FB231 reduced pathological α-synuclein and enhanced cell survival in human iPSC-derived dopaminergic neurons treated with α-synuclein preformed fibrils. Furthermore, FB231 attenuated α-synuclein pathology and dopaminergic neurodegeneration in a gut α-synuclein murine model of PD. Our findings support that Parkin plays a crucial role in maintaining neuronal homeostasis and that pharmacologic activation of Parkin may be a promising strategy to attenuate neurodegeneration in PD.

## Introduction

Parkin, an E3 ubiquitin ligase encoded by *PARK2*, is a central regulator of mitochondrial quality control and cellular homeostasis ^1,2^. As a key effector of the PINK1–Parkin mitophagy pathway, Parkin is recruited and activated when damaged mitochondria accumulate PINK1 on their outer membrane ^3,4^. Activated Parkin ubiquitinates mitochondrial surface proteins, targeting dysfunctional organelles for autophagic clearance—a process essential in neurons, where mitochondrial integrity is tightly linked to survival and neurodegeneration ^5–7^. Parkin also regulates apoptosis and the cell cycle machinery, and abnormal engagement of cell cycle progression may contribute to cell stress since these are critically regulated processes during development and terminal differentiation ^8,9^.

Mutations in *PARK2* are a leading cause of autosomal recessive juvenile Parkinson’s disease (PD). The most common mutations, including exon deletions and the recurrent R275W point mutation, result in loss of ubiquitin E3 ligase activity and defective mitophagy ^10,11^. The resulting accumulation of damaged mitochondria leads to increased oxidative stress and cellular toxicity ^12,13^, collectively contributing to the dopaminergic neurodegeneration characteristic of PD. *PARK2* mutations are highly penetrant and are associated with early-onset bradykinesia and tremors, a high incidence of lower-limb dystonia, and largely symmetric motor symptoms ^14^. Although PD patients with *PARK2* mutations respond to levodopa, they show a markedly higher rate of early treatment complications than those without *PARK2* mutations ^14^.

Dysregulation of Parkin function has also been implicated in sporadic PD ^15,16^. Oxidative stress and post-translational modifications ^17^, such as s-nitrosylation and phosphorylation, may impair Parkin enzymatic activity, resulting in a loss of Parkin function without a genetic mutation ^18^. Parkin is often insoluble or misfolded in the brains of patients with sporadic PD, suggesting that environmental and age-related factors may compromise its function ^19,20^. Furthermore, mitochondrial dysfunction and impaired mitophagy, hallmarks of sporadic PD, are closely linked to Parkin pathway dysregulation ^15^.

Current treatments for PD alleviate symptoms without halting disease progression. Given the pivotal role of Parkin in maintaining cellular health and in neurodegeneration, restoring its function represents a promising and potentially disease-modifying therapeutic strategy for PD. Several methods targeting the Parkin pathway are under investigation, including small molecules that activate Parkin, gene therapies to restore Parkin expression, and compounds that enhance PINK1 stability to promote Parkin recruitment ^21–23^.

A small-molecule compound, FB231, was recently reported to enhance Parkin activity through sensitizing cells to mitochondrial stress ^24^. Here, we examined FB231’s ability to activate Parkin E3 ligase activity. In addition, we tested the efficacy of FB231 in human iPSC-derived dopaminergic neurons treated with α-synuclein (αSyn) preformed fibrils (PFFs) and a murine gut αSyn PFF model. Our findings show that Parkin plays a crucial role in maintaining neuronal homeostasis and that pharmacologic activation of Parkin is neuroprotective against αSyn PFF toxicity.

## Results

### Parkin inactivation in neural stem cells disrupts cellular homeostasis and differentiation

To evaluate Parkin’s role in neural progenitor cells (NPCs), we generated Parkin knockout (KO) NPCs via CRISPR-Cas9 (Fig. 1a). Compared to wild-type (WT), Parkin KO NPCs exhibited disrupted neurosphere organization, irregular morphology, and increased aspect ratios (length/width ratio; Supplemental Fig. 1a–c), indicating impaired structural cohesion and cellular polarity. Further, proliferation was reduced in KO cells (Fig. 1b), accompanied by diminished BrdU incorporation (Supplemental Fig. 1d). Apoptosis was significantly elevated in Parkin-deficient NPCs, primarily upon etoposide challenge (Supplemental Fig. 1e), consistent with the role of Parkin in stress response and survival ^25^.

**Figure 1.**
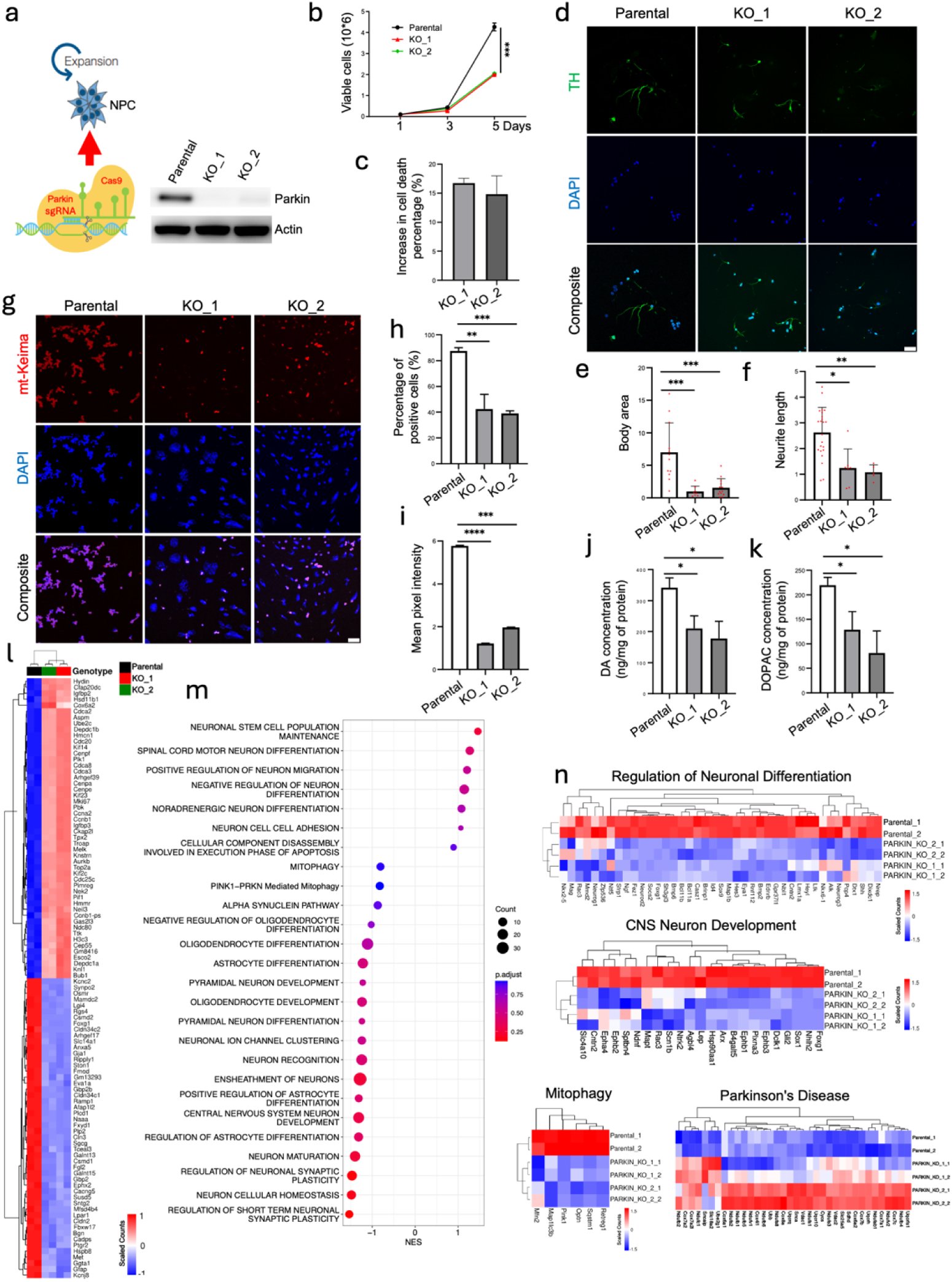
Parkin inactivation disrupts primary neural progenitor cell morphology and stability of differentiated neuronal cell state. (**a**) Diagram of the CRISPR-Cas9 system used to generate Parkin KO cells. Western blot was performed using cell lysates from parental NPC cells and two Parkin KO clones (i.e., KO_1 and KO_2), confirming loss of Parkin. (**b**) Parkin KO decreases the cell growth rate of neural progenitor cells. Red and green correspond to Parkin KO clones. Black corresponds to Parkin wild-type cells. (**c**) Parkin KO causes increased cell death upon differentiation of neural progenitor cells. Graph shows percent increase in cell death after 7 days in differentiation media. Error bars = 1 SD. (**d**) Immunofluorescence staining of tyrosine hydroxylase (marker of dopaminergic neurons) in differentiated Parkin KO and parental NPC cells. Scale bar = 50 µm. Quantitation showing changes in differentiated neurons related to average body area (**e**) and average neurite length (**f**). (**g**) Immunofluorescence assay of cells expressing mt-Kerma demonstrating that levels of mitophagy in parental NPC cells were more robust than in Parkin KO counterparts. Parkin KO deduces the percentage of mt-Keima positive cells (**h**) and fluorescence signal intensity (**i**). (**j**) HPLC (high-performance liquid chromatography) analysis of DA in differentiated Parkin KO and wild-type NPC cells. (**k**) HPLC analysis of DA-derived 3,4 dihydroxyphenylacetic acid (DOPAC) in differentiated Parkin KO and parental NPC cells. (**l**) Differentially expressed genes between Parkin wild-type and KO cells. Heatmap shows comparison of wild-type cell cultures (n=2, black) and Parkin KO cells (n=4) from KO_1 and KO_2 (red, green respectively; 2 replicates each). (**m**) GSEA analysis for neuron-related pathways using the ranked gene list from Parkin KO vs wild-type differentially expressed genes. Normalized enrichment scores (NES) show enrichment on the x-axis, while the adjusted p-value is shown using the color key on the right of the graph. Total genes in each gene set are represented as the size of the circle, as shown by the scale on the right. (**n**) Leading edge genes for GSEA analysis. Programs shown are for regulation of neuronal maturation, CNS neuron development, mitophagy, and PD pathways. Data presented as heatmaps, scaling each gene across samples with scales shown. Genes listed along the horizontal axis. Data from parental (wild-type) cells and Parkin KO clones are shown (2 clones each). All plots show individual data points, the mean, and SD. ANOVA and post-hoc Tukey’s test or student’s t-test (for f). *p<0.05, **p<0.01, ***p<0.001. ****p<0.0001.

Following induction of differentiation, Parkin KO NPCs exhibited increased cell death (Fig. 1c) and generated neurons and glia with abnormal morphologies. Tyrosine hydroxylase-positive (TH+) dopaminergic neurons exhibited smaller soma and shortened neurites (Fig. 1d–f), while GFAP+ astrocytes and RIP+ oligodendrocytes exhibited similar reductions in process length and cell body size (Supplemental Fig. 1f–k). These findings suggest that Parkin is required for stable neural differentiation across multiple lineages.

Next, we assessed mitophagy using mt-Keima, a pH-sensitive mitochondrial probe. WT NPCs displayed a strong mt-Keima red signal following FCCP/oligomycin treatment, indicating active mitophagy (Fig. 1g–i). In contrast, Parkin KO NPCs showed significantly reduced mt-Keima lysosomal localization, both in percentage of positive cells and red signal intensity, confirming impaired mitochondrial turnover ^26–28^. Lastly, we found that dopamine (DA) and its metabolite DOPAC were markedly reduced in differentiated Parkin KO neurons, indicating compromised dopaminergic function (Fig. 1j–k).

Together, these findings demonstrate that Parkin is essential for maintaining NPCs’ structural integrity, viability, and differentiation capacity, highlighting Parkin’s multifaceted role in early neural development and dopaminergic neuron specification.

### Parkin loss reshapes transcriptomes and disrupts programs required for neuronal homeostasis

Next, we examined the transcriptional alterations in NPCs after knocking out Parkin. Bulk RNA-seq of WT and KO NPCs showed extensive gene expression changes: 657 genes were significantly upregulated, while 1,440 were downregulated (Supplemental Fig. 2a). Among downregulated genes were Gja1, Anxa5, Gfap, and Osmr, regulators of mitochondrial function, cell survival, and inflammatory response ^29–33^ (Fig. 1l).

Gene set enrichment analysis (GSEA) revealed broad downregulation of pathways involved in neurogenesis, synaptic maintenance, and mitophagy, with compensatory upregulation of adhesion and stem-like transcriptional programs (Fig. 1m). Specific depletion of genes involved in neuron maturation (e.g., Neurod2, Sox1, Foxg1), CNS development (e.g., Ngf, Ntf5), and mitochondrial quality control (e.g., Pink1) was noted (Fig. 1n). Plotting the leading-edge genes demonstrated significant changes in the Kyoto Encyclopedia of Genes and Genomes (KEGG) PD pathway (Fig. 1n), ribosomal assembly (Supplemental Fig. 2b, c), and E2F/cell cycle regulation (Supplemental Fig. 2d).

DecoupleR-based transcription factor analysis showed increased activity of Olig1 and Olig2, and reduced activity of neurogenic and survival-promoting factors, including NeuroD2, Foxo3, Tp53, and Gata4 (Supplemental Fig. 2e) ^34^, suggesting a shift toward a proliferative, undifferentiated state.

These findings suggest that Parkin maintains transcriptional programs critical for neuronal maturation, mitophagy, and cell fate stability. Its loss skews NPCs toward proliferative, immature, and stress-vulnerable states, suggesting the inability of NPCs to properly differentiate into terminally differentiated neuronal cell types.

### Single-Cell Transcriptomics Reveal Cell Fate Disruption with Parkin Loss

To understand the impact of Parkin loss across distinct cell types, we performed single-cell RNA-seq of WT and KO differentiated cultures. Unsupervised clustering revealed four significant populations: neurons, astrocytes, oligodendrocytes, and proliferating cells (Fig. 2a; Supplemental Fig. 3a–d). Parkin KO cells showed increased oligodendrocytes, and proliferating cells (Fig. 2b), suggesting skewed fate trajectories.

**Figure 2.**
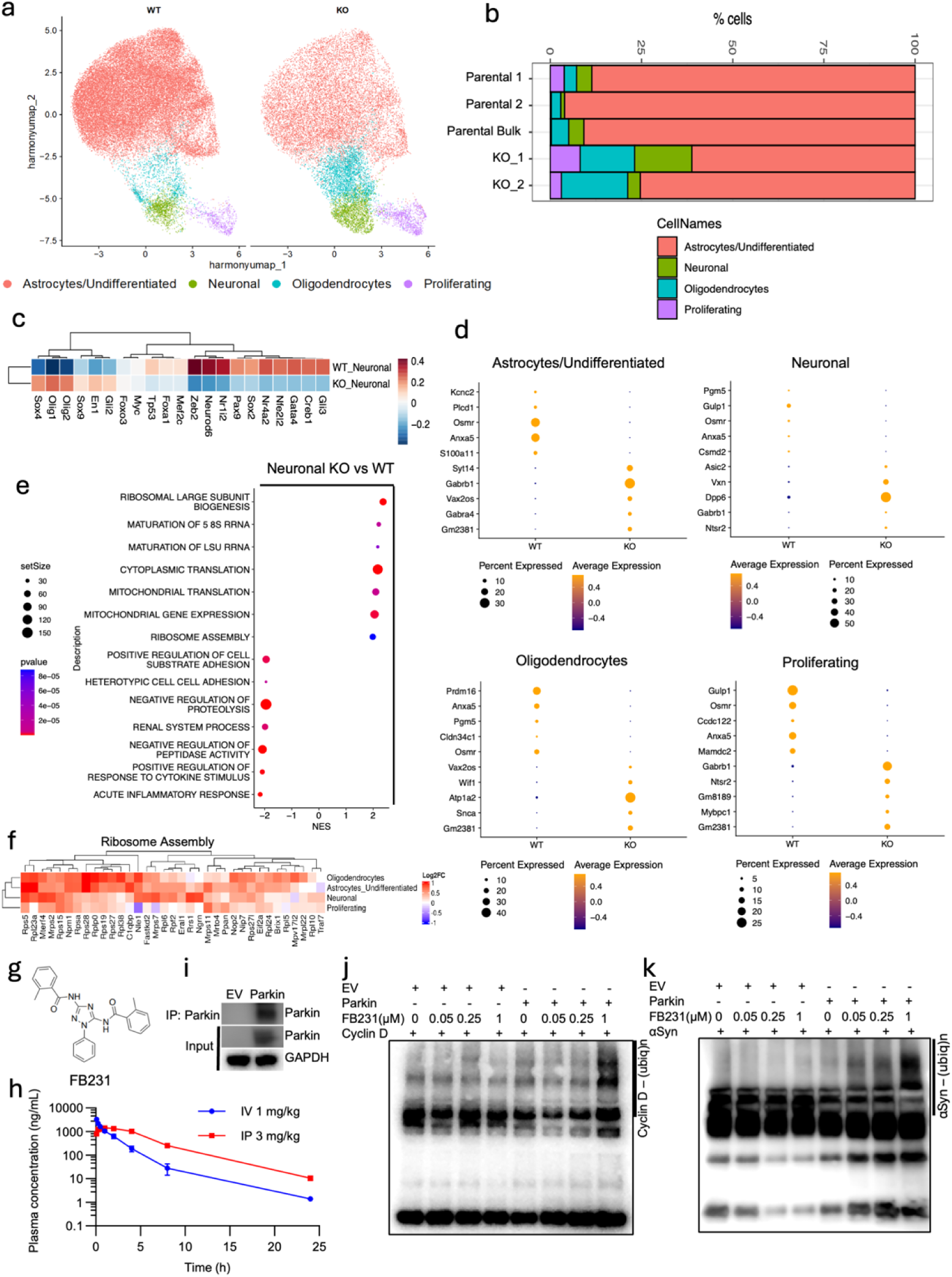
Parkin loss results in disrupted transcriptomes and lineage-specific gene expression programs. (**a**) Single cell sequencing results and unsupervised clustering of WT cells (Parental1, Parental2, Parental Bulk)(n=3) and KO cells (KO1, KO2)(n=2). Four primary clusters are identified that correspond to population types shown in the legend at the bottom. Colors indicate cell types. (**b**) Parkin loss alters the relative ratio of cell types following differentiation induction. Frequency of resulting cellular types from differentiation in each genotype is shown as a stacked bar plot for each sample. (**c**) Parkin KO results in alterations in transcription factors related to neuronal cell state. DecoupleR analysis was performed on differentially expressed genes between WT and KO neuronal-like cells. DecoupleR TF scores are plotted for each genotype. (**d**) Examples of Parkin KO-induced alterations in the expression of genes important for each cell type. Differential expression was performed to identify the top and bottom 5 genes by log2FC. Data shown as dot plots with the size of each dot representing the cell percentage expressing the gene, and the color scale indicating the average normalized expression level. (**e**) GSEA analysis of differentially expressed genes between KO neuronal cells and WT neuronal cells (x-axis represents the normalized enrichment score; dot size shows the gene set size; color shows p-value). (**f**) Genes from the gene ontology set Ribosome Assembly were selected, and the Log2FC is shown for KO vs WT cells as shown. Each row represents the KO vs WT comparison for the indicated cell type. (**g**) Chemical Structure of compound FB231. (**h**) Concentration of FB231 in the plasma of rats with IV and IP administration. Rats were treated with intravenous injection (1 mg/kg) and i.p. injection (3 mg/kg). (**i**) Immunoprecipitation (IP) of Parkin constructs. T98G cells were transfected with either pcDNA3.1 empty vector (EV) or with vector encoding WT Parkin. Cell lysates were prepared and immunoprecipitated with anti-Parkin antibody. (**j**) FB231 promotes Parkin activity to ubiquitinate cyclin D in vitro. Using Parkin IP, in vitro ubiquitination assay was performed. Different concentrations of compound FB231 were added in the indicated reactions. (**k**) Compound FB231 promotes Parkin activity to ubiquitinate αSyn in vitro. Using the above Parkin-pulled-down solution (Fig. 2i), an in vitro ubiquitination assay was performed, followed by a Western blot. Different concentrations of compound FB231 were added as indicated.

DecoupleR analysis of neuron-like cells revealed elevated expression of transcription factors associated with immature and glial fates (Olig1, Olig2) and reduced expression of neuronal regulators (Zeb2, Neurod6; Fig. 2c). Similar, though subtle, changes occurred in glial subsets (Supplemental Fig. 3e–f). TGFβ pathway activation was broadly elevated in KO cells, indicating stress-induced reprogramming (Supplemental Fig. 3g–i).

Differential gene expression across cell types revealed that Osmr and Anxa5 were consistently downregulated, particularly in astrocytic/undifferentiated cells, while Gabrb1 was upregulated in both neuronal and astrocytic populations (Fig. 2d). In oligodendrocytes, Prdm16, essential for mitochondrial biogenesis and neuronal migration, was also reduced ^35,36^. Genes related to membrane excitability (Syt14, Dpp6, Atp1a2) were increased ^37–39^.

Next, functional GSEA revealed increased translation and ribosome assembly programs in KO neurons and glia, alongside suppression of proteolysis and inflammation pathways (Fig. 2e–f; Supplemental Fig. 4a–c). Module scoring confirmed elevated ribosome/translation activity and diminished mitophagy signatures across all KO cell types (Supplemental Fig. 4d–f), consistent with heightened metabolic stress.

### FB231: A Parkin Activator with Favorable PK and Target Engagement

Given the neuroprotective role of Parkin, we aimed to utilize small-molecule activators of E3 ligases. Our recent publication includes the complete list of chemicals screened, along with the most favorable candidate, FB231 ^24^. First, we performed pharmacokinetic (PK) analysis of FB231 (Fig. 2g–h). We measured plasma FB231 concentrations using liquid chromatography-tandem mass spectrometry (LC/MS/MS) following a single intraperitoneal (i.p., 3 mg/kg) or intravenous (i.v., 1 mg/kg) injection (Fig. 2h) in rats. The half-life was 2.5 hours for i.v. administration and 3.2 hours for i.p. administration. FB231 remained detectable for up to 25 hours post-injection. Intraperitoneal administration resulted in high systemic exposure, with a systemic bioavailability of 75%. FB231 activated Parkin-dependent ubiquitination of both cyclin D1 and αSyn, two well-known targets of Parkin ^2,40^ (Fig. 2i–k).

### FB231 Attenuates ***α***Syn Pathology and Dopaminergic Neurodegeneration in a Gut ***α***Syn Mouse Model of PD

We tested the *in vivo* efficacy of FB231 in a gut-initiated αSyn pathology model of PD ^41,42^. First, we assessed FB231 levels in the brain and the plasma of mice. Following a 25 mg/kg i.p. injection in C57BL/6J mice, the brain-to-plasma ratios of FB231 were 2.3%, 1.5%, and 1.3% at 2-, 5-, and 24-hours post-injection, respectively, suggesting limited blood-brain barrier penetrance of the agent (Fig. 3a).

**Figure 3.**
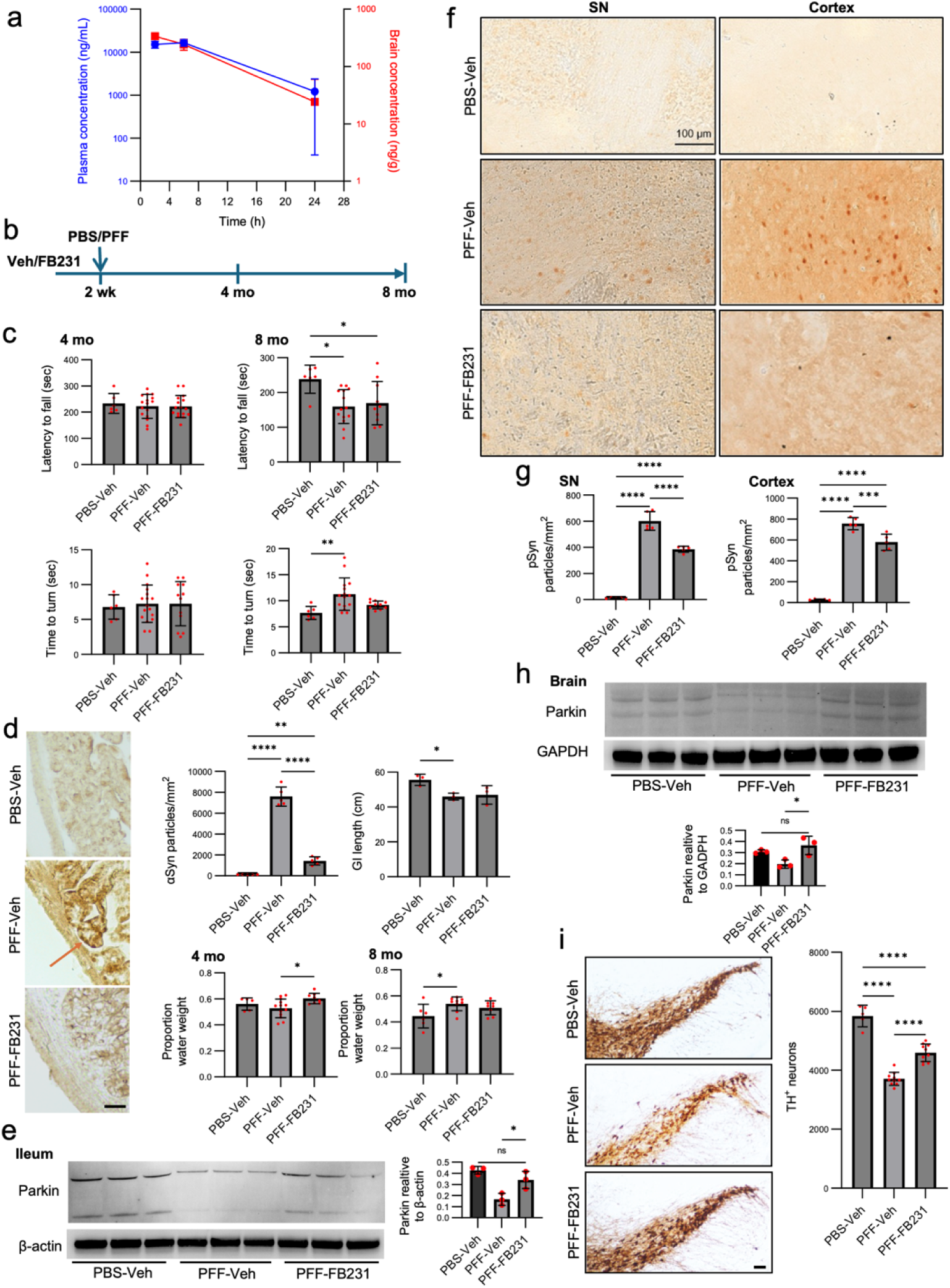
FB231 is neuroprotective in a gut αSyn mouse model of PD. (**a**) PK profile of FB231 in mice following systemic administration of FB231 (single injection i.p. at 25mg/kg). Plasma and brain FB231 concentrations were determined by LC/MS/MS. n=3 mice/group/time point. (**b**) Experimental schematic: C57BL/6J mice (male and female, 3-month-old) were pre-injected (i.p.) with FB231 25 mg/kg or Veh twice a week for 2 weeks. Mice then received a total of 25 µg PFF or PBS injection into 4 sites in the duodenal and pyloric muscularis layer. FB231 or Veh injection continued until the end of experiment at 8 months following PFF or PBS gut injection. (**c**) “Latency to fall” in the Rotarod test and “Time to turn” in the Pole test at 4 and 8 months after PFF gut injection. n=6-16 mice/group. (**d**) αSyn staining of the duodenum in αSyn in the duodenum of mice sacrificed 8 months post PFF injection, n=5/group. Arrow points to region of high αSyn expression. Scale bar = 50 µm. The staining was then quantified and analyzed across groups. Additionally, the GI length of mice sacrificed 8 months post-PFF injection was also quantified. Finally, the fecal water content of mice at 4 and 8 months after PFF injection. n=6-16 mice/group. (**e**) Parkin expression within ileum tissue at 8 months post-PFF injection was determined through immunoblotting, and Parkin signal was quantified relative to β-actin staining across different mice. n=3 mice/condition. (**f**) Immunostaining for pSyn in brain sections containing the SN and cortex at 8 months after PFF gut injection. (**g**) Quantification of pSyn particles presented in **e**. n=5 mice/group. (**h**) Parkin expression within brain tissue at 8 months post-PFF injection was determined through immunoblotting, and Parkin signal was quantified relative to β-actin staining. n=3 mice/condition. (**i**) Representative SN TH immunohistochemical sections collected at 8 months after PFF gut injection and stereological counting of TH-positive neurons. Scale bar = 150 µm. n=5-10 mice/group. All plots show individual data points, the mean, and SD. One-way ANOVA and Tukey’s post-hoc test was used for statistical analysis. *p<0.05, **p<0.01, ***p<0.001. ****p<0.0001. ns=not significant.

We employed a gastrointestinal (GI) injection model in which mice received either mouse αSyn PFFs (25 µg total) or PBS (phosphate-buffered saline) into the pyloric stomach and duodenum ^43^. Animals were pre-treated with FB231 or vehicle twice weekly, beginning two weeks before PFF administration and continuing until endpoint at 8 months (Fig. 3b). Body weight and general health were monitored throughout (Supplemental Fig. 5a). Two mice died in the PFF group and three in the PFF + FB231 group over the 8 months.

Motor function was assessed using the rotarod and pole tests ^44–46^. At 4 months, no significant differences were observed across groups. At 8 months, both PFF-Veh and PFF-FB231 groups displayed shorter latency to fall in the rotarod test, suggesting PFF-induced motor decline (Fig. 3c). The pole test revealed that PFF-Veh mice had significantly increased turn time compared to controls, while PFF-FB231 mice performed comparably to controls, indicating partial motor rescue by FB231.

We then examined GI pathology. Duodenal tissue from PFF-Veh mice showed prominent αSyn staining, which was reduced in PFF-FB231 mice (Fig. 3d). Quantification confirmed that FB231 significantly decreased PFF-induced αSyn accumulation in the GI tract. PFF-injected mice also exhibited shortened GI tract length, which was partially restored in FB231-treated mice (Fig. 3d; Supplemental Fig. 5b). Fecal water content, another measure of GI function ^47^, was elevated in PFF-Veh mice at 8 months, whereas FB231-treated mice maintained normal fecal water levels (Fig. 3d). The effect of FB231 on Parkin expression was assessed within ileum tissue (Fig. 3e). Quantification of the western blot revealed significantly lower levels of Parkin in PFF-Veh mice compared to the PBS-Veh group. Parkin expression was significantly higher in PFF-FB231 mice compared to PFF-Veh mice.

To assess brain pathology, we analyzed phosphorylated αSyn (pSyn) at Ser129, a hallmark of pathological aggregation ^48–50^. PFF-Veh mice displayed pSyn staining in both the substantia nigra (SN) and cortex, while PFF-FB231 mice exhibited significantly reduced pSyn accumulation in both regions (Fig. 3f–g). Control mice did not have detectable pSyn. Despite limited brain penetrance, FB231 appeared to increase Parkin expression in the brain tissue (Fig. 3h). Quantification revealed significantly higher expression of Parkin in PFF-FB231 mice than in controls.

We then evaluated dopaminergic neuron survival in the SN via immunostaining for TH. Both PFF-Veh and PFF-FB231 groups exhibited a loss of TH+ neurons relative to controls. However, FB231-treated animals had significantly more surviving TH+ neurons compared to vehicle-treated PFF mice, indicating a neuroprotective effect of FB231 (Fig. 3i).

Together, these data show systemic FB231 treatment reduces gut and brain αSyn pathology, preserves dopaminergic neurons, and improves motor and GI function in a gut-seeded PFF mouse model of PD.

### FB231 Enhances Mitophagy and Mitochondrial Integrity in Human iPSC-Derived Dopaminergic Neurons

To evaluate FB231 in a human-relevant system, we tested its effects in iPSC-derived dopaminergic neurons. We first confirmed dopaminergic neuron identity through expression of TH, dopamine transporter (DAT), and Tuj1 (Supplemental Fig. 6a). Treatment with FB231 (4–8 µM, 36 h) increased Parkin protein levels (Supplemental Fig. 6b), suggesting upregulation of Parkin-mediated pathways. To assess whether FB231 elicited any cellular stress responses, we compared our findings to Rosencrans, et al. ^24^, who reported mild mitochondrial toxicity and activation of the integrated stress response in proliferative cell lines in response to FB231 exposure. Western blot and qPCR analyses revealed that neurons treated with FB231 (4 µM) exhibited no significant induction of ATF-4, while ATF-3 was elevated compared to controls (Supplementary Fig. 6c–d). qPCR analysis of αSyn and Parkin revealed that FB231 exposure did not increase αSyn mRNA but elevated Parkin mRNA threefold (Supplementary Fig. 6e).

Live imaging with mitochondrial and lysosomal markers revealed enhanced colocalization of mitochondria with lysosomes after 24 h of treatment (Fig. 4a), indicating increased mitophagy. Compared to CCCP, which resulted in fragmented mitochondria due to depolarization ^51^, FB231 preserved elongated mitochondrial morphology. Morphometric quantification confirmed significantly higher levels of elongated mitochondria and colocalization in FB231-treated neurons compared to CCCP or DMSO (Fig. 4b; Supplemental Fig. 6f).

**Figure 4.**
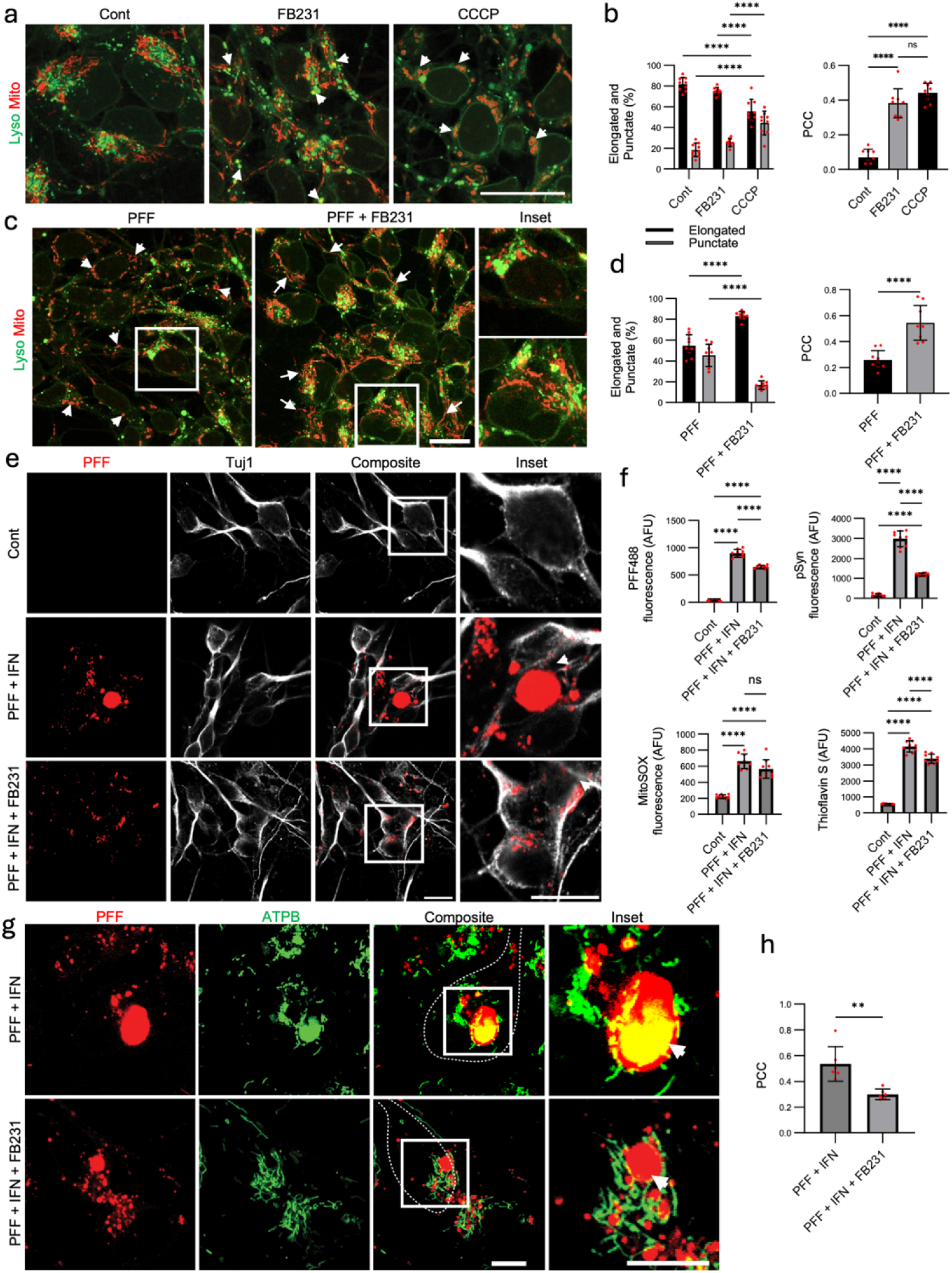
FB231 is neuroprotective in human iPSC-derived dopaminergic neurons. (**a**) Mitotracker (Mito) and Lysotracker (Lyso) staining and colocalization in dopaminergic neurons exposed to 4µM FB231, DMSO as a negative control, and 5µM CCCP as a positive control for 24 h. Arrowheads point to regions of mitochondrial colocalization with lysosomes. Scale bar = 20 µm. (**b**) Mitochondrial morphology, along with mitochondrial-lysosomal colocalization, was quantified across conditions using a custom Python script. Colocalization was quantified using the Pearson correlation coefficient (PCC). (**c**) Mito and Lyso staining and colocalization in neurons previously exposed to PFFs at 1 µg/mL for 24 h, then exposed to 4µM FB231 or DMSO for 24 h. Arrowheads point to punctate mitochondrial structures, while arrows point to elongated mitochondria. Scale bar = 20 µm. (**d**) Mitochondrial morphology, along with mitochondrial-lysosomal colocalization, was quantified across conditions using a custom Python script. (**e**) Differentiated dopaminergic neurons were exposed to the dual hit treatment. Arrowhead points to large PFF-positive punctate. Scale bar = 20 µm. (**f**) Unbiased quantification of PFF, pSyn, MitoSOX, and Thioflavin S across different conditions. (**g**) PFF-positive inclusions formed (arrowhead) in the presence and absence of FB231 exposure in neurons undergoing the dual-hit treatment at 14 days. Scale bar = 10 µm. (**h**) Colocalization of ATPB, a key component of mitochondrial ATP synthase, and PFF was calculated using a custom Python script. All plots show individual data points, the mean, and SD. ANOVA and post-hoc Tukey’s test or student’s t-test (for h) were used for analysis. *p<0.05, **p<0.01, ***p<0.001. ****p<0.0001. ns=not significant.

Next, we assessed FB231’s ability to counteract mitochondrial stress induced by PFFs (Fig. 4c–d; Supplemental Fig. 6g–h). iPSC-derived dopaminergic neurons were treated with human αSyn PFFs for 24 h, followed by FB231 or DMSO. PFF-exposed neurons exhibited punctate mitochondria, indicative of fragmentation ^52^, whereas FB231-treated neurons showed a marked restoration of elongated mitochondrial morphology. FB231 also significantly increased mitochondrial-lysosomal colocalization, reinforcing its ability to enhance mitophagy without causing mitochondrial toxicity, the latter being the mechanism by which CCCP induces/increases mitophagy.

### FB231 Protects Dopaminergic Neurons from ***α***Syn-Induced Stress and Inclusion Formation

To test neuroprotection under stress, we applied a dual-hit treatment using αSyn PFFs and interferon-gamma (IFN-γ), previously shown to induce Lewy body-like inclusions ^50^. After 10 days, neurons exposed to PFF + IFN-γ showed large PFF-induced positive αSyn inclusions and elevated pSyn, while FB231 treatment significantly reduced both PFF-induced αSyn burden and pSyn (Fig. 4e–f; Supplemental Fig. 7a–b). FB231 also reduced Thioflavin S fluorescence, a marker of β-sheet-rich aggregates, and partially decreased mitochondrial superoxide levels (MitoSOX), though the latter did not reach statistical significance (Fig. 4f; Supplemental Fig. 7c). In neurons treated with PFFs alone (no IFN-γ), FB231 significantly reduced both MitoSOX and PFF signal, demonstrating its protective effect under milder stress conditions (Supplemental Fig. 7d).

### FB231 Preserves Neuronal Morphology and Prevents Mitochondrial Incorporation into Inclusions

Next, we assessed the structural integrity of dopaminergic neurons, focusing on indicators of overall neuronal health. In PFF + IFN-γ-treated cultures, neurons showed neurite degeneration, clustering, and TH/Tuj1 signal loss (Supplemental Fig. 7e–f). FB231 partially restored neuron number, neurite length, and morphology, suggesting preservation of neuronal health and architecture. On a transcriptional level, FB231 increased Parkin mRNA levels while reducing αSyn expression, when comparing PFF+FB231-treated cells to the PFF-only condition (Supplementary Fig. 7g)

Finally, we examined whether FB231 reduced mitochondrial sequestration into PFF-induced αSyn inclusions. Our previous work showed that dysfunctional mitochondria accumulate inside these Lewy body-like inclusions ^50^. Immunostaining for ATPB revealed significantly lower colocalization between mitochondria and αSyn aggregates in FB231-treated neurons compared to untreated PFF + IFN-γ samples (Fig. 4g–h), indicating preservation of mitochondrial function and exclusion from aggregates.

## Discussion

Our findings demonstrate that Parkin is essential for maintaining neuronal cell state and preserving dopaminergic neuronal phenotypes. Parkin directly regulates key genes and pathways governing neuronal maturation and stem-like cell transitions. Notably, we identified dysregulation of several PD-associated molecular pathways, mitophagy, and processes required for homeostasis in Parkin-deficient neurons. We characterized a small-molecule Parkin activator, FB231, that enhances Parkin E3 ligase activity. Finally, we demonstrated that FB231 treatment reduces pathological αSyn accumulation and promotes dopaminergic neuronal survival in human iPSC-derived dopaminergic neurons and a mouse PD model of gut-to-brain αSyn transmission.

We provide evidence that Parkin loss impairs neuronal differentiation and the maintenance of neuronal cell states. The observed disruptions in neurite stability and morphology suggest that Parkin regulates neuronal structural integrity, potentially through its ubiquitin ligase activity. Gene expression analyses further indicated that Parkin modulates pathways critical for neuronal maturation and stability, supporting a model in which Parkin activity is necessary for proper neuronal lineage specification. These findings align with prior studies linking Parkin function to neuronal plasticity and survival ^53–55^. The severe morphological and functional consequences of Parkin loss highlight the need for Parkin activators to restore Parkin function and prevent or modify neurodegeneration in PD.

Among the small Parkin activators we developed, compound FB231 ^24^, is well-tolerated in rodents, with minimal adverse effects on vital organs, behavior, body weight, or survival. Consistent with a previous report by Rosencrans, et al. ^24^, the ability of FB231 to enhance Parkin activity in our model systems is supported by increased protein expression, increased mitophagy markers, and ubiquitination of known targets. The favorable profile of FB231 highlights the feasibility of testing the compound’s efficacy in preclinical models.

Previous studies have linked Parkin and αSyn in cellular-ubiquitinated inclusions through their interaction with an αSyn-interacting protein, and αSyn is an established target of Parkin ^56^. Parkin deficiency results in neurodegeneration and accumulation of pathological pSyn in monkeys, and Parkin expression reduces pSyn aggregates in a rat PD model of αSyn overexpression ^54,57^. In our gut αSyn PFF-injected mice modeling “body-first” PD ^41^, chronic FB231 treatment appeared to provide sustained Parkin activation in the gut and, unexpectedly, in the brain despite minimal blood-brain barrier penetrance. FB231-treated mice exhibited improved performance in the pole test at a later time point. GI abnormalities, including αSyn accumulation, were partially restored in FB231-treated mice. In the brain, FB231 treatment reduced αSyn pathology in both SN and cortex and TH-positive neuronal loss in the SN. It is to be determined whether the neuroprotective effect of FB231 in the gut model arises from direct action of FB231within the brain, or from systemic mechanisms that reduce pathological αSyn before its transmission to the brain, or a combination of both. Further studies are needed to delineate the specific site(s) of FB231 action and determine where it exerts the greatest neuroprotective effect.

In human iPSC-derived dopaminergic neurons exposed to αSyn PFFs and an inflammatory stressor, FB231 treatment significantly reduced αSyn pathology and mitochondrial oxidative stress. Importantly, FB231 prevented neurite degeneration and maintained dopaminergic neuronal integrity. Although exposure to FB231 did not prevent the formation of Lewy body-like inclusions, it prevented healthy mitochondria from being incorporated within these inclusions. Unlike CCCP, which induced mitophagy through mitochondrial toxicity in iPSC-derived dopaminergic neurons, FB231 enhanced mitophagy without inducing overt mitochondrial damage, as treated neurons maintained elongated mitochondrial morphology.

The upregulation of ATF-3 observed in our dopaminergic neuronal model was overall consistent with the previous study’s findings ^24^; however, we detected no induction of ATF-4 and no reduction in cell viability following FB231 treatment. This discrepancy may reflect differences in dosing regimens and exposure duration, but more likely arises from intrinsic cell-type–specific factors such as metabolic demand, mitochondrial turnover, and the threshold for stress signaling. Unlike proliferating tumorigenic cells that depend heavily on glycolysis and are particularly vulnerable to mitochondrial perturbation ^58^, terminally differentiated dopaminergic neurons rely primarily on oxidative metabolism and maintain robust antioxidant and quality-control mechanisms ^59^. These features may enable neurons to tolerate enhanced Parkin activity without fully engaging ATF-3/4-mediated ISR pathways. Overall, our findings highlight the importance of evaluating candidate compounds in disease-relevant cellular and animal models, as FB231 exhibits distinct effects in neurons versus tumor-derived cell lines. Furthermore, our experiments were conducted under endogenous Parkin expression, suggesting that overexpression systems may exaggerate certain responses not observed in physiologic contexts. Future studies should delineate the downstream molecular mechanisms underlying FB231-mediated neuroprotection in neuronal systems.

In conclusion, our study identifies Parkin as a central regulator of neural differentiation and homeostasis and provides preclinical evidence that small-molecule Parkin activation is a viable strategy to mitigate αSyn pathology and dopaminergic neurodegeneration. These findings lay the groundwork for developing Parkin-targeting small molecules as a promising avenue for disease-modifying interventions in PD and potentially other neurodegenerative disorders characterized by mitochondrial dysfunction and αSyn aggregation.

## Methods

### Cell culturing

#### mNPC Cell culture and differentiation

Murine neural progenitor cells were a gift from Dr. Jason Huse ^60^. The primary mNPCs were maintained in NeuroCult Basal Medium containing NeuroCult Proliferation Supplement, 20 ng/ml EGF, 10 ng/ml basic FGF, 2 μg/ml heparin (Stemcell Technologies), 50 units/ml penicillin, and 50 μg/ml streptomycin (Thermo Fisher Scientific). Cells were cultured in a humidified incubator at 5% CO2 and 37 °C. To induce differentiation, cells were plated on dishes coated with 10 μg/ ml laminin (Sigma-Aldrich) and cultured in NeuroCult Basal Medium containing NeuroCult Differentiation Supplement (Stemcell Technologies). For experiments, cells were used at passages 5–10.

#### iPSC differentiation and culturing

Human iPSC-Derived Neural Progenitor Cells (STEMCELL, Catalog # 200-0620) were expanded and cultured in Neural Progenitor Medium (STEMCELL, Catalog # 05833). Following two passages, cells underwent midbrain differentiation using the Midbrain Neuron Differentiation Kit (STEMCELL, Catalog # 100-0038) supplemented with 200 ng/mL of Human Recombinant Shh (STEMCELL, Catalog # 78065) for 7 days. Dopaminergic NPCs were then plated on 6/24/96-well plates for Western blotting, fluorescent imaging, and plate-reader quantification, respectively. NPCs were then matured into neurons using the Midbrain Neuron Maturation Kit (STEMCELL, Catalog # 100-0041) for 2 weeks. Following maturation, cells were maintained in BrainPhys Neuronal Medium and SM1 Kit (STEMCELL, Catalog # 05792).

### Construction of Parkin KO mNPC cell line using CRISPR/Cas9

CRISPR-Cas9-nickase editing was performed according to a published protocol^61^. pSpCas9n(BB)-2A-GFP (PX461) was a gift from Feng Zhang (Addgene plasmid # 48140). Briefly, optimize guide RNAs (GAGTGGTTGCTAAGCGACAGG and GTGTCAGAATCGACCTCCACT) were designed and cloned into the PX461 plasmid. Transient transfection was performed using the Nucleofector™ Kit for Mouse Neural Stem Cells (Lonza) according to the manufacturer’s guidelines. Individual GFP-positive cells were sorted into single wells containing culturing medium. Single-cell-derived colonies were gradually expanded and screened for frameshift mutations in all alleles using targeted PCR of the locus followed by TOPO-TA cloning (Invitrogen) and sequencing of individual colonies. Two independent mNPC Parkin KO cell clones were picked to perform the experiments.

### Cell proliferation assay with flow-based trypan blue cell viability analyzer

Mouse mNPC cells were grown in triplicate on laminin-coated (10 μg/ml) 6-well plates (1 x 105 cells per well) and stopped at different time points. Proliferation was evaluated using the flow-based trypan blue cell viability analyzer (Vi-Cell XR, Beckman Coulter).

### Annexin V-FITC apoptosis assay

The presence of apoptosis of cells was detected using the Annexin V-FITC Apoptosis Detection Kit (Clontech Laboratories, Inc.) according to the manufacturer’s protocol. Briefly, cells were treated with 100 μM Etoposide for 18 h, harvested, rinsed, and resuspended with Binding Buffer, and then incubated with Annexin V and Propidium Iodide at room temperature in the dark. Flow cytometry was performed using FACSymphony A3 Flow Cytometer (BD Biosciences). Ten thousand events were acquired for each sample. Results were quantified using FlowJo software.

### BrdU incorporation assay

The BrdU detection kits were purchased from BD Biosciences to examine BrdU that has become incorporated into a cell’s DNA when it is going through cell division. Cells were stained and analyzed using FACSymphony A3 Flow Cytometer (BD Biosciences). Ten thousand events were acquired for each sample. Results were quantified using FlowJo software.

### Western blotting

#### For NPC experiments

Cells were lysed in cell lysis buffer (Invitrogen) containing protease inhibitor cocktails.

Loading sample was prepared after protein estimation. Equal amounts of protein were loaded per lane and separated by 4-12 % SDS-PAGE. Proteins in the gels were transferred to the polyvinylidene difluoride membrane, followed by blocking with 5 % (w/v) skim milk in TBST. Then, the membranes were incubated with respective primary antibodies, anti-Parkin (Cell Signaling, CS4211) or anti-β-actin (Sigma, A3853 or A2066) overnight at 4°C, followed by incubation with HRP-conjugated secondary antibodies (1:3,000) for one hour at room temperature. Chemiluminescence reagent ECL was used to visualize the protein bands (Amersham ImageQuant 800).

#### Immunoprecipitation

Cells were harvested and lysed in lysis buffer (50 mM Tris, pH 7.4, 250 mM NaCl, 5 mM EDTA, 50 mM NaF, 1 mM Na_3_VO_4_, 1% Nonidet P40 (NP40), 0.02% NaN_3_) containing protease inhibitors. Lysates were clarified by centrifugation at 12,000 × g for 10 min, and protein concentration was determined using a BCA assay kit (Thermo). For immunoprecipitation assays on endogenous proteins, samples were incubated with antibodies (10 μl of anti-PARKIN (4211, Cell Signaling); or 3.3 μl non-specific mouse IgG (Invitrogen)). Proteins were precipitated by using protein A-sepharose beads that had been blocked with 3% powdered milk. The beads were washed four times with lysis buffer and aliquoted for in vitro ubiquitination assay.

#### Mouse and iPSC samples

Differentiated neurons, cultured on 6-well plates, were lifted with Accutase (STEMCELL, 07920), pelleted, and stored at -80°C. Lysates were prepared using the RIPA Lysis Buffer System (Santa Cruz, sc-24948A). Protein concentration was then analyzed for each sample using the Pierce BCA Protein Assay Kits (Thermo Scientific, 23225). Samples were then heated for 15 min at 75°C in NuPAGE LDS Sample Buffer (Invitrogen, NP0007). Lysates were blotted together on the same gel (Invitrogen, NP0322BOX).

Membranes were incubated with primary antibodies overnight at 4°C following blocking with 5% Bovine Serum Albumin (Millipore Sigma, A9647-100G) in TBS with Tween (Thermo Scientific Chemicals, J77500.K2) for 1 h. Primary antibody incubation was done overnight, while secondary antibody incubation was done at room temperature for 1 h. Membranes were imaged using ChemiDoc (Bio-Rad) and analyzed using Image Lab software (Bio-Rad).

For blot quantification, the signal intensity of the antibody of interest (i.e., Parkin; Invitrogen, 740019R) was divided by the signal intensity of the control GAPDH (Sigma-Aldrich, AB2302). The signal intensity was quantified with ImageJ (NIH).

### Quantitative Reverse Transcription PCR (qRT-PCR)

Total RNA was extracted using TRIzol reagent (Invitrogen, 15596026) per manufacturer’s instructions. Cells were lysed in TRIzol, mixed with chloroform, and centrifuged to isolate the aqueous phase. RNA was precipitated with isopropanol, washed with 70% ethanol, air-dried, and resuspended in RNase-free water. Purity and concentration were assessed by NanoDrop (A260/A280 ≥ 1.8). For cDNA synthesis, 1 µg RNA was reverse transcribed in a 20 µL reaction using PrimeScript RT Master Mix (Takara, RR036A; 37 °C 15 min, 85 °C 5 s, hold 4 °C). cDNA was diluted fivefold and stored at −20 °C. qPCR was performed in triplicate using SYBR Green Universal PCR Master Mix (Applied Biosystems, 4309155) on a StepOnePlus system (Applied Biosystems). Each 10 µL reaction contained 5 µL SYBR mix, 1 µL cDNA, 0.5 µL each primer (200 nM), and 3 µL water. Cycling: 50 °C 2 min, 95 °C 2 min, 40 cycles of 95 °C 15 s/60 °C 1 min, with a melt curve. Relative expression was calculated by 2^−ΔΔCT using housekeeping genes as controls.

Forward and Reverse Primers for ATF-3 and ATF-4 were obtained from Origene (HP233668 and HP205494). ATF-3’s forward primer sequence was CGCTGGAATCAGTCACTGTCAG, and reverse primer sequence was CTTGTTTCGGCACTTTGCAGCTG. ATF-4’s forward primer sequence was TTCTCCAGCGACAAGGCTAAGG, and reverse primer sequence was CTCCAACATCCAATCTGTCCCG. GADPH primers (Origene, HP205798) had the following sequence for forward and reverse primers, respectively: GTCTCCTCTGACTTCAACAGCG, ACCACCCTGTTGCTGTAGCCAA. Parkin primers (Origene, HP207857) had the following sequence for forward and reverse primers, respectively: CCAGAGGAAAGTCACCTGCGAA and CTGAGGCTTCAAATACGGCACTG. Finally, αSyn primers (Origene, HP200326) had the following sequence for forward and reverse primers, respectively: ACCAAACAGGGTGTGGCAGAAG and CTTGCTCTTTGGTCTTCTCAGCC.

### Immunohistochemical, immunofluorescent staining, and confocal microscopy

#### NPC experiments

Samples were fixed in 4% PFA for 25 minutes RT and permeabilized with PBS 0.1% Triton-X100 for 3 minutes on ice, blocked with PBS 3% BSA, 0.3% Triton-X100 for 2 hours RT, followed by overnight incubation with primary antibody at 4° C (0.5 µg/ml rabbit Tomm20 antibody FL-145; Santa Cruz Biotechnology) diluted in PBS 0.1% BSA, 0.3% Triton-X100. The secondary goat anti-rabbit antibody conjugated with DyLight 649 (Jackson ImmunoResearch) was applied for 1hour at room temperature at a concentration of 2.8µg/ml in conjunction with 1 µg/ml Hoechst33342. Cells were imaged using an Olympus ScanR automated microscope equipped with a motorised stage and 20x APO planar objective. 18 images were acquired for each well using the following combination of excitation/emission filters: Hoechst33342 was excited through a 350/50 nm band pass filter, and fluorescence intensity was collected through a 460/30 band pass filter. YFP was excited through a 500/20 nm band pass filter, and fluorescence intensity was collected through a 540/35 band pass filter. DyLight 649 was excited through a 640/30 nm band pass filter, and fluorescence intensity was collected through a 700/75 band pass filter. Images were processed and analysed as described in the Image Analysis section.

Other antibodies used were GFAP monoclonal 1:500 (Abcam, ab33922); FoxA2 monoclonal 1:300 (Abcam, ab108396); Tyrosine Hydroxylase (TH) monoclonal 1:10,000 (ImmunoSTAR 22941); and RIP monoclonal 1:200 (Cell Signaling, CS3493). After washing, the cells were fluorescently labeled with secondary antibodies anti-rabbit IgG (H+L), F(ab’)2 Fragment (Alexa Fluor® 488 Conjugate) #4412 (Cell Signaling, CS4412) or anti-mouse IgG (H+L), F(ab’) 2 Fragment (Alexa Fluor® 555 Conjugate) (Cell Signaling, CS4409). After counterstaining with 4,6-diamidino-2-phenylindole (DAPI, Vector Labs), the cells were observed under the confocal microscope (Leica Microsystems).

#### Confocal imaging of mitophagy with mito-Keima

Parental or PARKIN KO mNPC cells stably expressing mito-Keima were treated with oligomycin (5 µM) + Antimycin (5 µM) for 18 hr. Mito-Keima, excited at 561 nm, was artificially colored as red under the confocal fluorescent microscope. 4,6-diamidino-2-phenylindole (DAPI, Vector Labs) was used for cell counterstaining.

#### Live imaging: mitophagy

Live imaging of dopaminergic neurons was done following the plating of dopaminergic NPCs on glass-bottomed plates (MatTek, P35G-1.5-14-C) and maturing them into dopaminergic neurons over the course of 2 weeks. Prior to imaging, cells then underwent PFF-only, PFF+IFN-γ, and/or FB231 (or DMSO) exposure. On the day of imaging, neurons were stained with Mitotracker and Lysotracker (Invitrogen, L7526 and M7510) for 30 min before imaging them on a Nikon C2 Confocal Microscope.

#### Mouse brain tissue

Mice were sacrificed, and their brains were collected. Brains were post-fixed and cryoprotected. Brains were sectioned coronally, and serial sections were collected as described ^62,63^. Sections were incubated with anti-pSyn at Ser129 (p-syn/81A, BioLegend, 825701) at 1:500 at 4°C overnight and then incubated with secondary antibody for 1 h at 37 °C. The staining was developed by incubating with DAB. Quantification of pSyn staining was performed using the optical fractionator method at 40× magnification (Olympus BX51 microscope and Olympus CAST stereology software) to count positively stained particles in sections containing different brain regions ^64^.

A complete set of serial midbrain sections were collected and immunostained for TH using anti-TH antibody (clone TH2, Sigma, T1299) at 1:1000 at 4°C overnight. Sections were counterstained for Nissl. Unbiased stereological counting was performed blindly as previously described ^62,63^. The number of TH-positive neurons was counted across sections, and the total number of cells per individual animal was then calculated.

#### Immunofluorescence imaging of dopaminergic neurons

For each biological replicate, neurons from separate differentiation events were plated and matured on glass coverslips (Thomas Scientific, 1217N79), previously treated with Matrigel (Corning, 356234) in 12 or 24-well plates (Thermo Scientific, Nunc). Cells underwent the experimental procedure and were fixed using 4% PFA (Thermo Scientific Chemicals, J61899AP).

All coverslips were washed with PBS and stained for pSyn (Abcam, ab51253), Tuj1 (Abcam, ab41489), and TH (Genetex, GTX113016), following permeabilization and blocking with 5% Bovine Serum Albumin (Millipore Sigma, A9647-100G) and 0.05% of 10% Triton (Thermo Scientific Chemicals, A16046.AE). Secondary antibodies tagged with Alexa 488 and Alexa 568 (Invitrogen, A-11008 and A-11031) were used following overnight incubation with primary antibodies. Finally, prior to mounting with Prolong Diamond (Invitrogen, P36961), cells were stained with DAPI (Thermo Scientific, 62248). Fluorescent imaging was done using a Nikon C2 Confocal Microscope and viewed using NIS-Elements software (Nikon).

### Detection of DA and DOPAC

DA and DOPAC were assessed by HPLC-ECD (high-performance liquid chromatography electrochemical detection) using published protocols ^65–67^. Cell pellets were resuspended in PE buffer (0.1 mM EDTA, 1 μM DHBA (internal standard), 40 mM phosphoric acid) at a ratio of 50 μL PE buffer per 1 million cells. The suspension was freeze-thawed for 3 cycles, then centrifuged at 14,000 rpm for 15 minutes at 4°C. For brain tissue samples, 200 μL PE buffer was added, and tissues were homogenized. The supernatant was then transferred to a 0.22 μM Co-star spin column and centrifuged at 14,000 rpm for 5 minutes at 4°C. The resulting filtrate was run through an Ultimate 3000 UHPLC system with a Zorbax Reversed-Phase C18 Column (150 mm x 3.0 mm i.d., 5 μm particles). An isocratic mobile phase was used to achieve separation of the analytes of interest. The mobile phase contained 75 mM sodium phosphate, 1.7 mM sodium 1-octosulfonate, 100 μL/L triethylamine, 5 μM EDTA, and 10% acetonitrile, and was adjusted to a final pH of 4.2 using 85% phosphoric acid. A flow rate of 0.6 mL/min was used for a run time of 17 minutes per sample. The sample injection volume was 10 μL. ECD was performed using an analytical cell with two channels set at -150 mA and 250 mA. The DA and DOPAC contents were calculated as ng per mg wet tissue for brain samples and ng per mg protein for cell samples.

### In vitro ubiquitination assay

In vitro ubiquitination assays were performed using a commercial ubiquitination kit (Enzo Life Sciences Inc.) per the manufacturer’s protocol. Different concentrations of compounds were added to the reactions. After incubation for 30-60 min at 37°C, the reactions were terminated and analyzed with Western blotting.

### Compounds

FB231 was synthesized by Wuxi App Therapeutics. A detailed synthesis schema has been reported ^24^.

### Image analysis

#### NPC data

Images were processed and analyzed using Columbus HCS Analysis software (Version 2.5.0, PerkinElmer) as follows: Tomm20 fluorescence intensity was corrected using the parabola algorithm. Hoechst 33342 fluorescence was used to identify and count cells. Cells were segmented according to Tomm20 fluorescence intensity. Spot detection was optimized to recognize the number and total cellular area of Tomm20-stained clusters (mitochondria). Tomm20 staining intensity, spot numbers, and spot area were used to train a linear classifier algorithm that discriminated between Tomm20-positive (high intensity, spot numbers, and spot area) and Tomm20-negative cells (low intensity, spot numbers, and spot area). Bar graphs were generated reporting the number of Tomm20 negative cells expressed as percentage of total cells imaged for each well. Results were shown as mean ± SD of 3 experiments performed in triplicate.

#### Quantification of mitochondrial morphology and organelle colocalization

A custom script was developed to detect and analyze mitochondrial and lysosomal movement, morphology, and colocalization in time-lapse images of dopaminergic neuronal cultures. The code, written in Python, is available at: https://doi.org/10.5281/zenodo.15490490. This allowed for automatic, unbiased quantification of the imaging dataset.

For quantification of colocalization, we used Pearson correlation coefficient (PCC). PCC evaluates the relationship between pixel intensity values in two channels (e.g., mitochondria and lysosomes), providing a measure of how consistently the intensities of one channel vary with those of the other across pixels. A high positive PCC indicates that when one fluorophore exhibits high intensity at a given location, the other does as well, suggesting spatial coincidence, molecular interaction, and functional association. Conversely, a value near zero implies random spatial distribution, and a negative PCC indicates mutually exclusive localization.

#### Neurite length

Neurites were traced using ImageJ (NIH), and their lengths were quantified using the NeuronJ plugin for ImageJ, as described by the plugin’s developers, Meijering, et al. ^68^. The results were then analyzed using Prism (GraphPad).

#### Fluorescent quantification via plate reader

All fluorescent quantification (except for Neurite Length) was done using a FluoSTAR Omega (BMG LABTECH) plate reader for unbiased quantification. Cells were fixed or imaged live, depending on the assay/experiment. For each quantification experiment, the proper excitation and emission filters were selected prior to quantification. Bottom optics were used, and Spiral sampling was selected, allowing for the detection of fluorescence from multiple different areas of each well.

For quantification of MitoSOX (Invitrogen, M36008), cells were washed and incubated with PBS prior to the addition of MitoSOX reagent. All biological replicates were then imaged consecutively on separate plates.

For antibody staining, samples were fixed separately but stained simultaneously using the protocol described above.

### PFF production and characterization

#### In vivo experiments

Recombinant mouse αSyn was purified as described ^69^. Endotoxin was removed using the Pierce high-capacity endotoxin removal resin (Thermofisher, 88270) and levels quantified using a Pierce chromogenic endotoxin kit (Thermofisher, A39552S). Endotoxin levels were 0.06 EU/µg protein for mouse alpha-synuclein and 0.09 EU/µg protein for human alpha-synuclein. PFFs were generated by diluting momoner to 5 mg/mL in 150 mM KCl, 50 mM Tris-HCl (pH 7.5), shaking for seven days at 37°C. PFFs were then spun for 10 min, resuspended in buffer, and a small volume dissociated in guanidinium chloride (8M), and protein concentration was measured using A280. The fibrils were then brought to a final concentration of 5 mg/mL, aliquoted into 22 µL aliquots, snap frozen, and stored at -80°C. Fragments were generated using a sonicator (QSonica, Q700) with a 16°C water chiller and size confirmed using dynamic light scattering (Wyatt technologies, Dynapro Nanostar).

#### In vitro experiments

αSyn PFFs were prepared as previously described ^50,70,71^. Fibril size was verified prior to and after sonication (∼81 nm) using Dynamic Light Scattering. Following characterization, PFFs used in dopaminergic neurons were tagged with fluorescent tags, as described by Maneca, et al.^72^.

### Experimental animals

For animal experiments, FB231 was suspended in Veh containing 10% DMA, 15% Solutol, and 75% 20%HPBCD and administered i.p., alternating sides of the abdomen. For experiments in iPSCs, FB231 was suspended in DMSO.

For the PK study in SD rats, animals were injected with FB231 either i.p. (3 mg/kg) or i.v. (1 mg/kg). Plasma samples were collected at multiple time points post-injection. To determine concentrations of FB231 in plasma vs. brain tissue, C57BL/6 mice were injected with FB231 i.p. at 25 mg/kg. Plasma and whole brain tissue were collected at 2, 6, and 24 h.

To test the efficacy of FB231, 3-month-old male C57Bl/6 mice were purchased from the Jackson Laboratory. Mice were maintained in home cages at a constant temperature with a 12-h light/dark cycle and free access to food and water. All animal protocols were approved by the Massachusetts General Hospital Animal Care and Use Committee (#2018N00003).

### LC/MS/MS determination of FB231 in biological samples

Plasma (10 µL) and brain (50 µL) samples were prepared using protein precipitation with an internal standard (tolbutamide in methanol/acetonitrile). Brain tissue was first homogenized in MeOH:PBS (3:7, 1:4 w/v) using a high-speed dispersator and stored at −80 °C. Both plasma and brain samples were vortexed, centrifuged, and the resulting supernatants were used for LC/MS/MS analysis. The method employed linear regression with 1/(x²) weighting, with LLOQs of 10 ng/mL for plasma and 0.1 ng/mL for brain homogenate. LC/MS/MS analysis was performed using an AB Sciex Triple Quad 6500+ with electrospray ionization in negative mode and MRM detection. FB231 (410.10 → 158.90) and tolbutamide (268.80 → 169.80) were monitored. Samples were separated on an Atlantis Premier BEH C18 AX column using a water/acetonitrile gradient with 5 mM NH Ac and 0.05% FA. The column temperature was 40 °C, and the injection volume was 4 µL. Separate gradients were used for brain and plasma samples. PK parameters, including half-life and bioavailability, were calculated from plasma concentration-time profiles.

### Intestinal intramuscular **α**Syn PFF injection

Adapted from a previously established protocol ^41^, mice were anesthetized with isoflurane, and a midline abdominal incision was made. The pyloric stomach and upper duodenum were gently exposed. Using a 10 µL Hamilton syringe with a 33 GA needle, αSyn PFF was injected into the wall of the pyloric stomach (2 sites, 6.25 µg/2.5 µl per site) for a total of 12.5 µg αSyn PFF, and into the intestinal wall of the duodenum (2 sites, 6.25 µg/2.5 µl per site) for a total of 12.5 µg αSyn PFF. Control mice were injected with an equivalent volume of PBS at the same sites. Mice were returned to their housing.

### Rotarod test

The rotarod test was performed to evaluate motor coordination and balance as previously described ^64^. Mice were placed on a fixed speed (16 rpm) rotating rod (3.0 cm) (Rotamex, Columbus Instruments). Three trials were performed, and the greatest value for each session was used for analysis. The time mice spent on the rotating rod was calculated up to a maximum of 240 seconds.

### Pole test

The pole test was performed to test motor coordination and motor abnormalities as previously described ^73^. Three trials were performed. Time taken for the mice to turn downward was recorded.

### Fecal water

GI function tests will include colon motility, which will be monitored by a one-hour stool frequency assay ^74^. The total stools are weighed to provide a wet weight, then dried overnight at 65 °C and weighed again to provide a dry weight. Water content is calculated as ((wet weight-dry weight)/wet weight)x100).

### Measurement of GI length

Following euthanasia in accordance with institutional ethical guidelines, the entire gastrointestinal tract was harvested from each mouse by carefully dissecting from the stomach to the rectum. The small intestine and colon were separated at the cecum. Intestinal contents were flushed with ice-cold phosphate-buffered saline (PBS) using a 20 mL syringe fitted with a 20G plastic catheter. Each segment was then opened longitudinally using fine scissors and gently spread flat. Tissues were laid out without tension on a clean surface, and total lengths of the small intestine and colon were measured using a calibrated ruler. Care was taken to avoid stretching or compression of the tissue during measurement.

### PFF treatment of dopaminergic neurons

We adapted the treatment regimen conducted previously by Bayati, et al. ^50^. Briefly, mature dopaminergic neurons were exposed to αSyn PFFs at a 1 µg/mL concentration for 48 h. Following fresh media change, cells were incubated for 72 h, prior to exposure to IFN-γ (Gibco, 300-02-500UG) for 48 h. Depending on their randomly assigned condition, cells were then either treated with DMSO or with 4 µM of FB231 for 36 h. Neurons were then incubated in fresh media for 36 h prior to fixation/collection for imaging, quantification, or Western blotting. For most experiments, neurons underwent a total of 10 days in culture following initial treatment with PFFs.

### Statistical analysis

All data are presented as mean ± standard deviation (SD). Statistical analyses were performed using GraphPad Prism version 10.3.1. For comparisons between two groups, unpaired two-tailed Student’s t-tests were used. For comparisons involving more than two groups, one-way or two-way analysis of variance (ANOVA) was applied, followed by Tukey’s post hoc test. Biological replication was ensured through the use of independently derived neural progenitor or iPSC-dopaminergic neuronal cultures, and *in vivo* experiments were conducted using independent cohorts of mice. For imaging and functional assays, technical replicates were averaged within each biological replicate prior to statistical testing. For single-cell RNA sequencing and bulk transcriptomic analyses, differential expression was evaluated using adjusted p-values with false discovery rate (FDR) correction, and pathway enrichment significance was determined using GSEA. Sample sizes and exact statistical tests for each experiment are detailed in the corresponding figure legends. p-values ≤0.05 were considered statistically significant for all analyses.

## List of Abbreviations

αSyn: Alpha-synuclein
ANXA5: Annexin A5
ATF-3: Activating transcription factor 3
ATF-4: Activating transcription factor 4
ATP1A2: Adenosine triphosphatase (Na+/K+) transporting, alpha 2 polypeptide
ATPB: ATP synthase subunit beta
BCA: Bicinchoninic acid assay
BSA: Bovine serum albumin
BrdU: Bromodeoxyuridine
cDNA: Complementary DNA
CCCP: Carbonyl cyanide m-chlorophenyl hydrazone
CNS: Central nervous system
CRISPR: Clustered regularly interspaced short palindromic repeats
DA: Dopamine
DAB: Diaminobenzidine
DAT: Dopamine transporter
DAPI: 4′,6-diamidino-2-phenylindole
DLS: Dynamic light scattering
DMA: Dimethylacetamide
DMSO: Dimethyl sulfoxide
DOPAC: 3,4-dihydroxyphenylacetic acid
DPP6: Dipeptidyl peptidase-like protein 6
EDTA: Ethylenediaminetetraacetic acid
ECD: Electrochemical detection
EGF: Epidermal growth factor
EV: Empty vector
FGF: Fibroblast growth factor
FITC: Fluorescein isothiocyanate
FOXA2: Forkhead box A2
FOXO3: Forkhead box O3
Gabrb1: Gamma-aminobutyric acid receptor subunit beta-1
GAPDH: Glyceraldehyde-3-phosphate dehydrogenase
GATA4: GATA binding protein 6
GFAP: Glial fibrillary acidic protein
GI: Gastrointestinal
GJA1: Gap junction protein alpha 1
GOBP: Gene ontology biological process
GSEA: Gene set enrichment analysis
HPLC: High-performance liquid chromatography
HRP: Horseradish peroxidase
IFN-γ: Interferon-gamma
i.p.: Intraperitoneal
i.v.: Intravenous
iPSC: Induced pluripotent stem cell
IP: Immunoprecipitation
ISR: Integrated stress response
KEGG: Kyoto Encyclopedia of Genes and Genomes
KO: Knockout
LC/MS/MS: Liquid chromatography–tandem mass spectrometry
LLOQ: Lower limit of quantification
Lyso: Lysotracker dye
Mito: Mitotracker dye
MitoSOX: Mitochondrial superoxide indicator
mNPC: Murine neural progenitor cell
NGF: Nerve growth factor
NEUROD2: Neuronal differentiation 2
NEUROD6: Neuronal differentiation 6
NPC: Neural progenitor cell
NP-40: Nonidet P-40
NTF5: Neurotrophin 5
Olig1: Oligodendrocyte transcription factor 1
Olig2: Oligodendrocyte transcription factor 2
OSMR: Oncostatin-m receptor
PFA: Paraformaldehyde
PFF / PFFs: Preformed fibrils (αSyn)
PFF633: Alexa Fluor 633–labeled PFFs
PCC: Pearson correlation coefficient
PD: Parkinson’s disease
PINK1: PTEN-induced kinase 1
PK: Pharmacokinetics
PI: Propidium iodide
PRDM16: Pr/set domain 16 protein
PVDF: Polyvinylidene fluoride
qPCR: Quantitative polymerase chain reaction
qRT-PCR: Quantitative reverse-transcription pcr
RIPA: Radioimmunoprecipitation assay
RIP: Receptor-interacting protein (oligodendrocyte marker)
RNA-seq: RNA sequencing
RT: Room temperature
SDS-PAGE: Sodium dodecyl sulfate–polyacrylamide gel electrophoresis
Shh: Sonic hedgehog
SN: Substantia nigra
Sox1: SRY-related HMG-box 1
STAT1/3: Signal transducer and activator of transcription 1/3
SYT14: Synaptotagmin 14
TBS: Tris-buffered saline
TBST: Tris-buffered saline with tween-20
TEM: Transmission electron microscopy
TF: Transcription factor
ThT: Thioflavin T
TH: Tyrosine hydroxylase
TOMM20: Translocase of the outer mitochondrial membrane 20
TP53: Tumor protein p53
UMAP: Uniform Manifold Approximation and Projection
Veh: Vehicle
WT: Wild type
YFP: Yellow fluorescent protein
Zeb2: Zinc finger e-box binding homeobox 2

## Author Contribution

X.C. and T.A.C. conceived and supervised the study; Y.G., A.B., T.J.A., F.Z., J.A.J., T.A.C., and X.C. designed the experiments; Y.G., A.B., F.Z., Y.Z., C.S., and J.A.J. performed in vitro and in vivo studies; A.F., S.N.G., L.A.V., W.L., T.M.D., S.D., and M.A.S. provided critical reagents and conceptual advice; Y.G., A.B., T.J.A., P.P., F.Z., V.M., and J.G.S. carried out data processing, bioinformatic, and statistical analyses; Y.G., A.B., T.J.A., J.A.J, T.A.C, and X.C. interpreted the data and wrote the manuscript.

## Funding

This research was funded in part by Aligning Science Across Parkinson’s No. ASAP-000312 through the Michael J. Fox Foundation for Parkinson’s Research (MJFF) and by the National Institute of Health through the National Institute of Neurological Disorders and Stroke grant R01NS102735.

## Materials & Correspondence

Materials and correspondence should be addressed to Corresponding authors: Xiqun Chen and Timothy A. Chan at.

## Competing interests

J.A.J. is an employee of Snohamer LLC, the parent company of Finsno Bio; serves as the CEO of NysnoBio, of which Snohamer is also the parent company; has received funding from Michael J. Fox Foundation for aspects of this work; and is an inventor on a patent describing FB231 (US10308617B2). T.A.C. acknowledges grant funding from Bristol-Myers Squibb, AstraZeneca, Illumina, Pfizer, An2H, and Eisai. T.A.C. has served as an advisor for Bristol-Myers, MedImmune, Squibb, Illumina, Eisai, AstraZeneca, and An2H. T.A.C. holds ownership of intellectual property on using tumor mutation burden to predict immunotherapy response, with a pending patent, which has been licensed to PGDx. All other authors claim no competing interests.

## Data Availability

All data supporting the findings of this study are included within the article and its Supplementary Information. Bulk RNA-sequencing and single-cell RNA-sequencing datasets generated in this study will be deposited in the NCBI Gene Expression Omnibus (GEO) and will be publicly available upon publication (accession codes to be provided). The custom Python code used for automated quantification of mitochondrial and lysosomal morphology, motility, and colocalization is publicly available via **Zenodo** at https://doi.org/10.5281/zenodo.15490490

Source data underlying all figures are available from the corresponding authors upon reasonable request. No restrictions apply to the availability of the code or publicly deposited datasets. Access to any additional raw imaging data not included in the Supplementary Information may be subject to reasonable request due to file size and storage constraints.

## Supplementary Figure Legends

**Supplementary Figure 1.**
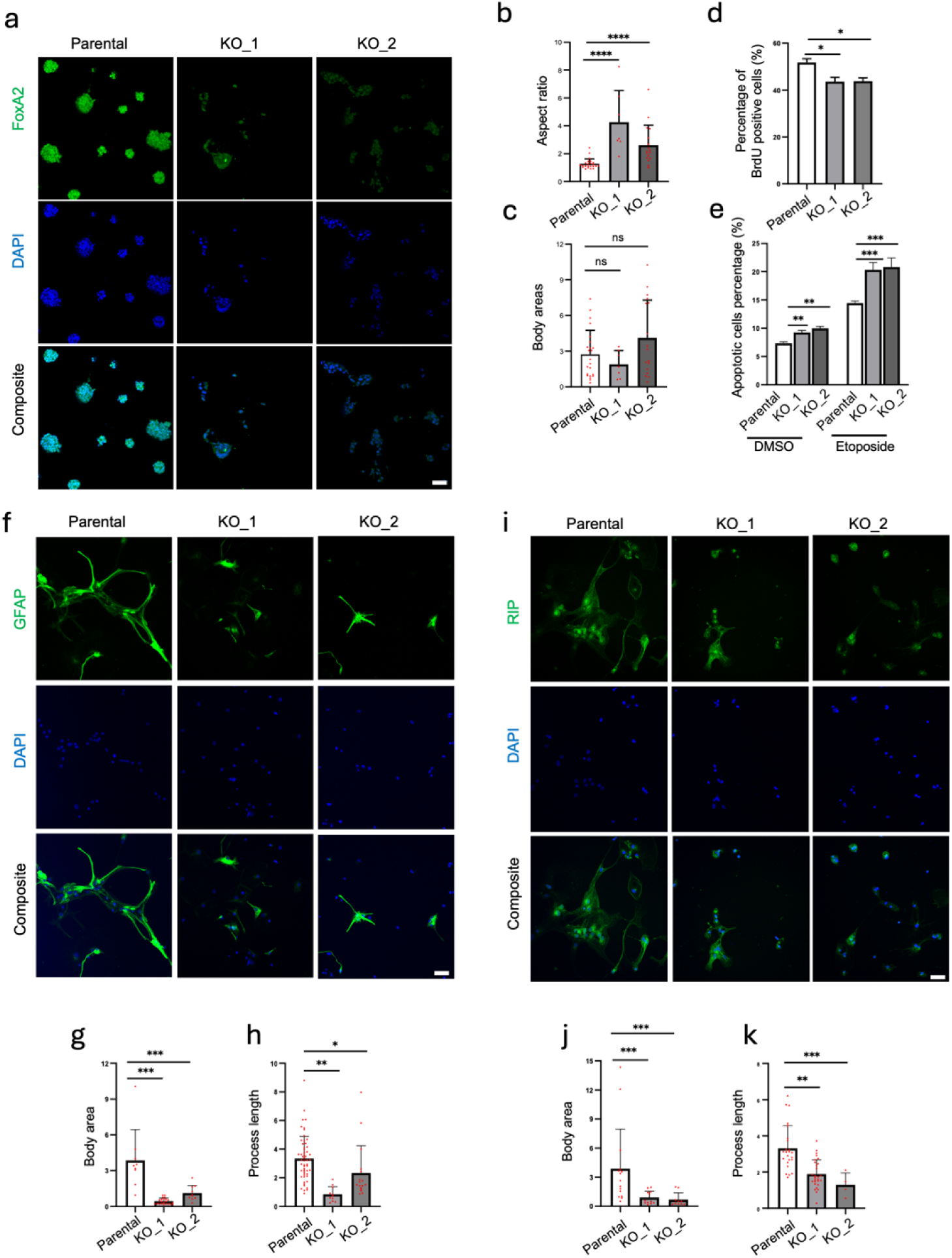
Parkin KO results in phenotypic abnormalities in NP cells. (**a**) Immunofluorescence staining of FoxA2 in Parkin KO and parental (WT) NPC cells. Parkin KO disrupts the normal shape and structure of cell bodies and cell aggregation. Scale bar = 50 µm. (**b**) Quantitation of average aspect ratios and (**c**) cell body sizes. (**d**) Flow cytometry shows Parkin KO NPCs have a decreased number of cells in S phase. (**e**) Flow cytometry shows Parkin KO NPC cells have an increase of apoptotic cells both at baseline and after etoposide treatment. (**f**) Immunofluorescence staining of GFAP (marker of astrocytes) in differentiated Parkin KO and parental cells. Quantification of astrocyte phenotypes showing (**g**) difference in body area between the genotypes and (**h**) differences in average process length. (**i**) Immunofluorescence staining of RIP (marker of oligodendrocytes) in differentiated Parkin KO and parental cells. Quantitation of oligodendrocyte phenotypes showing **(j**) difference in body area between the genotypes and (**k**) differences in average process length. All plots show individual data points, the mean, and SD. ANOVA and post-hoc Tukey’s test or student’s t-test (for f). *p<0.05, **p<0.01, ***p<0.001. ****p<0.0001. ns=not significant.

**Supplementary Figure 2.**
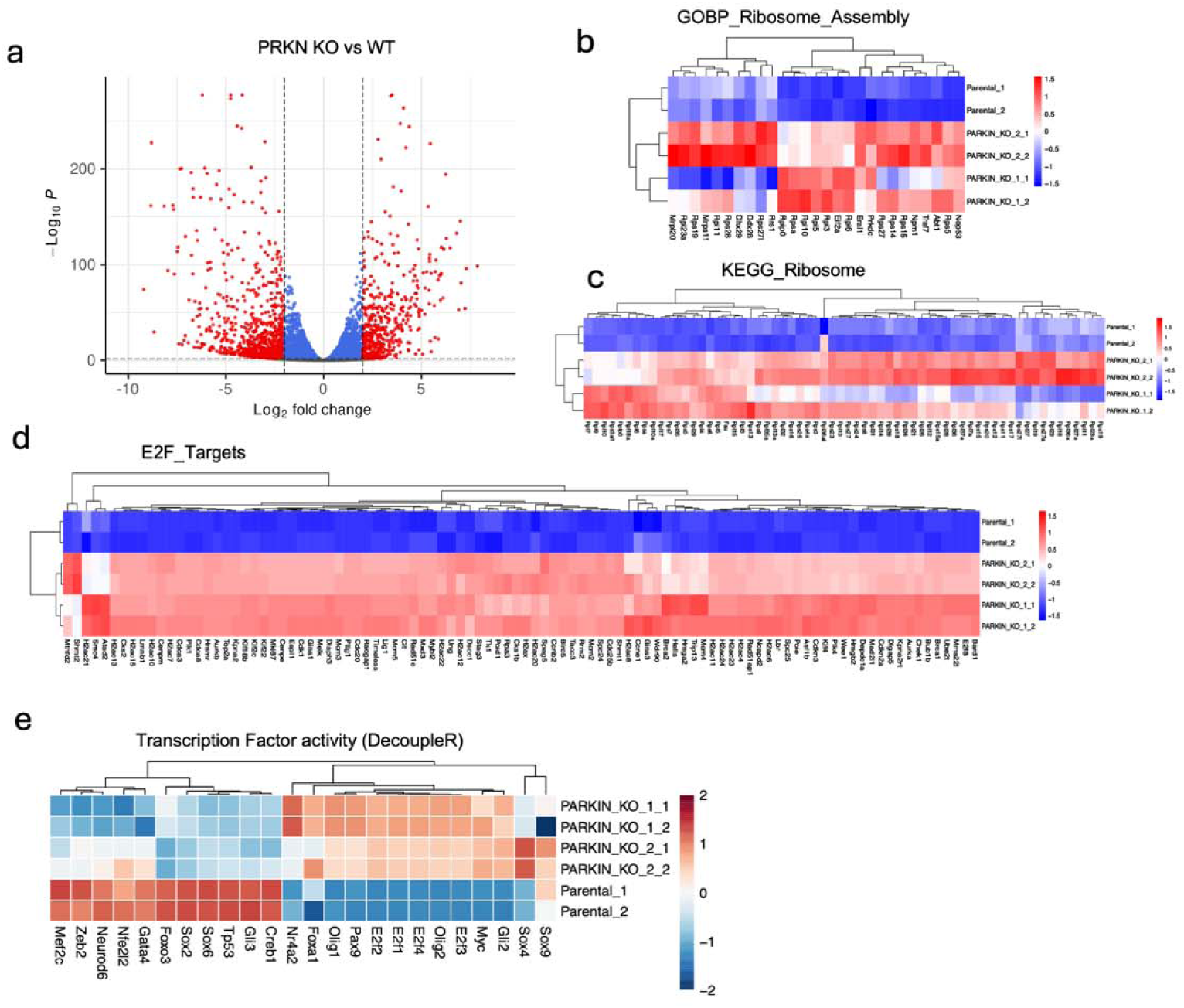
Parkin KO-induced changes in the transcriptome. (**a**) Gene expression changes in Parkin KO neural progenitor cells. Volcano plot showing differentially expressed genes comparing KO (n=4) and WT (n=2) samples. Crosshairs show cutoffs for adjusted p-values <0.05 and Log2FC >2 or <-2. Red dots represent those values which met our threshold of p-adjusted values <0.05 and Log2FC >2 or <-2; blue and grey dots indicate not significant. (**b-d**) Examples of gene expression programs undergoing large-scale changes due to Parkin inactivation. GSEA analysis of differentially expressed genes between KO vs WT cells showing leading edge genes for pathways of GOBP Ribosome assembly, KEGG Ribosome, and E2F targets. Data are plotted in heatmaps with gene values scaled across samples. (**e**) Differential transcription factor activity in Parkin KO versus WT cells. DecoupleR was used to identify pathway activity. Pathways with top differential activity scores are shown.

**Supplementary Figure 3.**
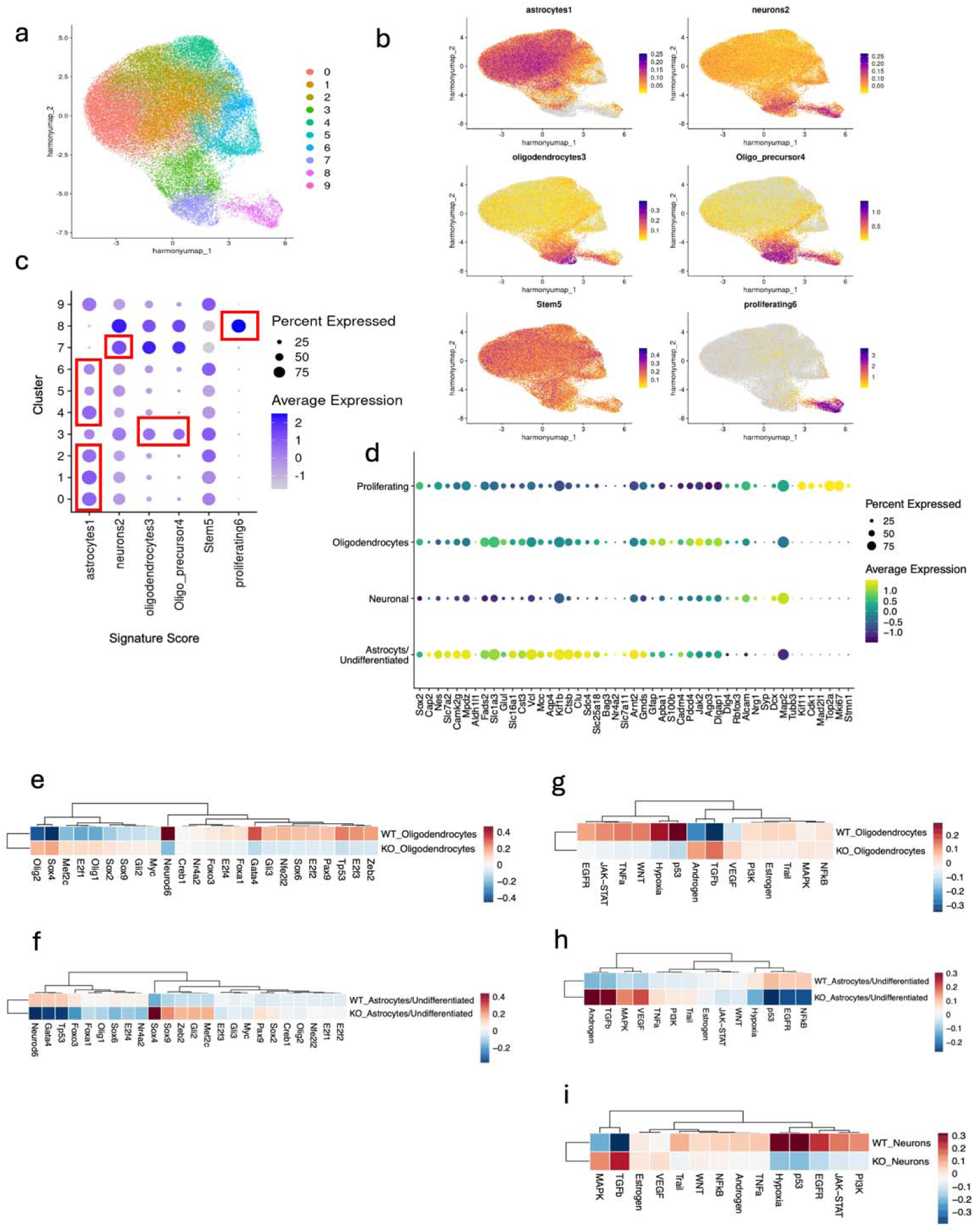
Single-cell sequencing of differentiated Parkin KO neural and glial cells reveals lineage-specific gene expression defects. (**a**) UMAP showing clusters identified in differentiated cells. Unsupervised clustering was performed using Seurat, with Harmony batch correction to generate a single harmonized UMAP with 10 clusters. n=2 KO and n=3 WT Parkin clones. (**b**) Gene set overlays identify astrocytes, neurons/neuron-like cells, oligodendrocytes, oligodendrocyte precursors, stem cells, and proliferating cells. Data generated and scored using Seurat’s AddModuleScore function. Gene set scores are shown as feature plots, highlighted where the gene set of interest is most highly enriched on the UMAP. (**c**) Four primary classes of differentiated cells were identified and related to their corresponding clusters: astrocytes, neuronal, oligodendrocytes, and proliferating (marked with red boxes). (**d**) Key cell lineage markers across primary classes of differentiated cells. After naming the 4 cellular classes by grouping clusters into similarly scored groups, key genes distinguishing each class are shown by gene. For each dot, expression level is represented by the color scale and size of each dot represents the percentage of cells expressing a given gene. (**e, f**) Altered neurodevelopmental-related transcription factor activity in Parkin KO cells. DecoupleR transcription factor analysis of Parkin KO vs WT oligodendrocytes and astrocytes/undifferentiated cells. (**g-i**) DecoupleR pathway activity analysis of Parkin KO vs WT cells. Neuronal, oligodendrocyte, and astrocytes/undifferentiated cells are shown, highlighting the top-most altered pathways.

**Supplementary Figure 4.**
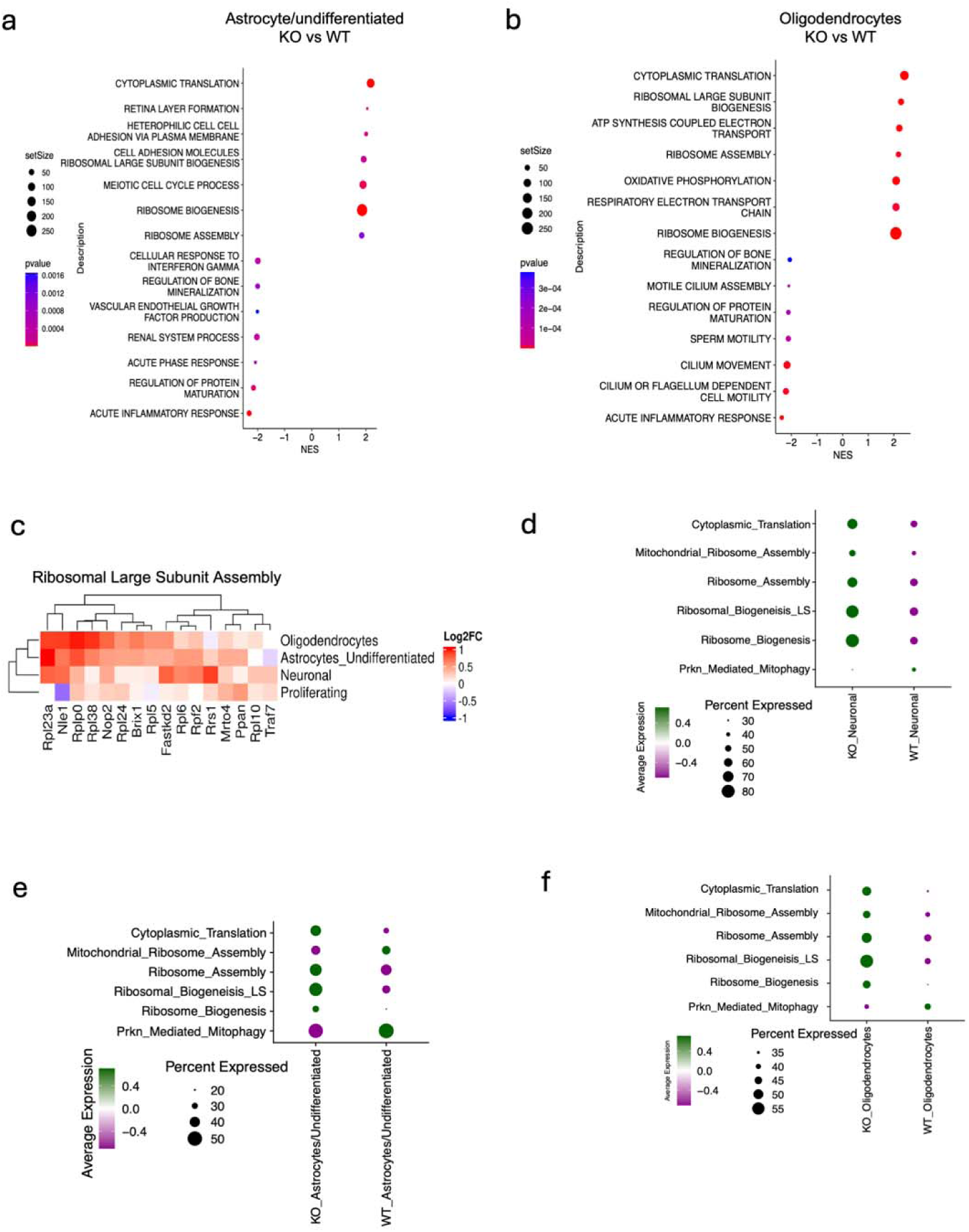
Gene expression programs altered in Parkin KO astrocytes and oligodendrocytes. (**a**) Differential gene expression of Parkin KO vs WT astrocyte/undifferentiated cells. Data shows output of GSEA analysis using the GOBP gene set. Most enriched pathways are shown. NES, normalized enrichment score. Dot size represents the input gene set size, and color scale shows the p-value. (**b**) Differential gene expression of Parkin KO vs oligodendrocytes. Data shows output of GSEA analysis using the GOBP gene set. Most enriched pathways shown. NES, normalized enrichment score. Dot size represents the input gene set size, and color scale shows the p-value. (**c**) High expression of ribosome assembly genes in Parkin KO cells. Differential expression was performed for each cell type, and the log2FC value of genes from the ribosomal large subunit assembly pathway from GOBP were plotted. Color scale shows the log2FC value of each comparison. (**d-f**) Relative expression of gene sets for ribosomal pathways, translation, and mitophagy are shown. Seurat’s AddModuleScore was used to score individual cells across the entire dataset for each gene set. After scoring cells, the labelled cell types for each genotype were plotted, where dot features indicate the average expression score of each gene set and the percent of cells expressing the gene set.

**Supplementary Figure 5.**
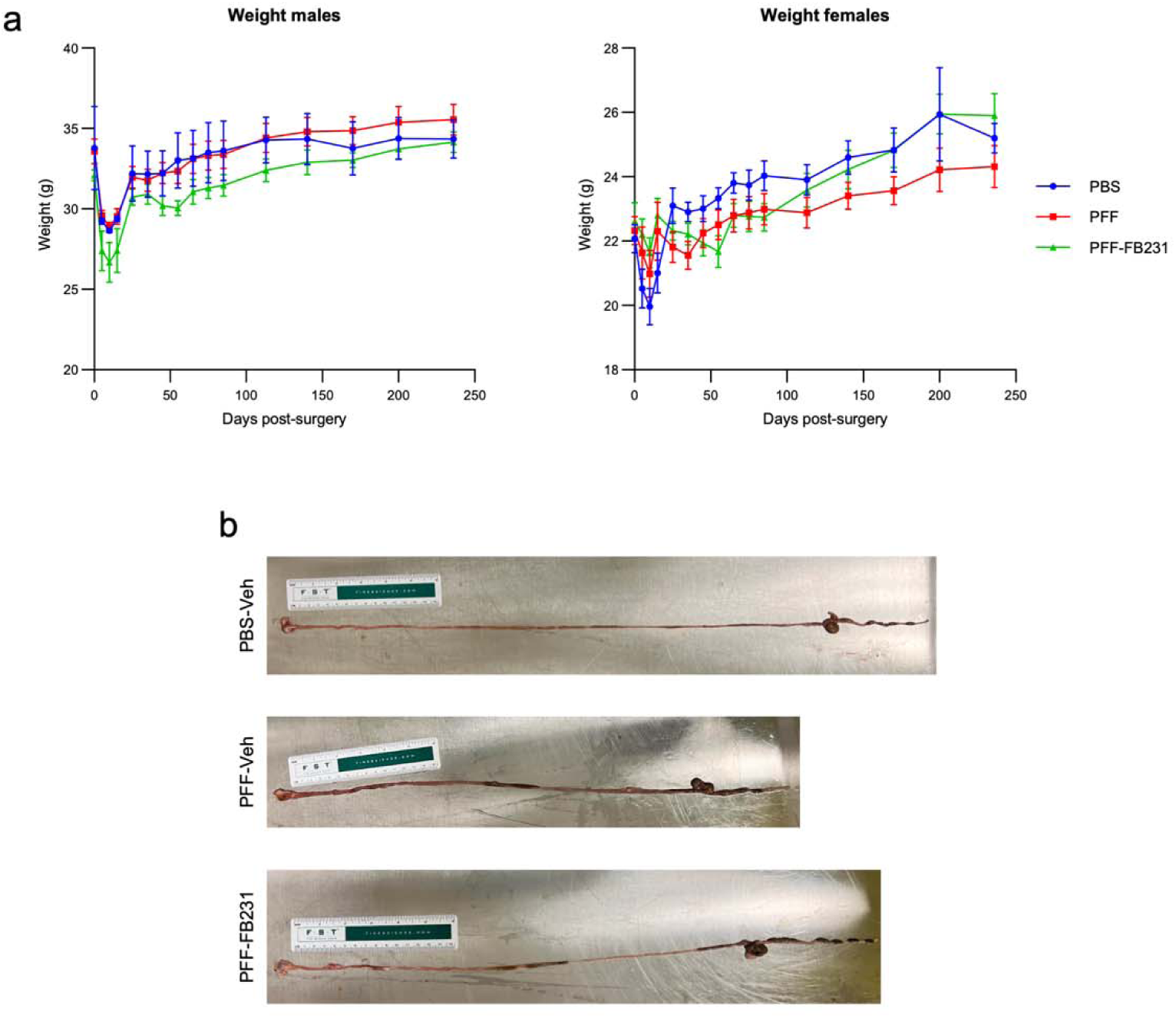
Mice body weight and GI length measurement. (a) The body weight of the mice used in our experiments was monitored over 8 months across the different experimental conditions for both male and female mice. n=6-16 mice/group. (b) Representative GI length measurement of mice in different experimental groups.

**Supplementary Figure 6.**
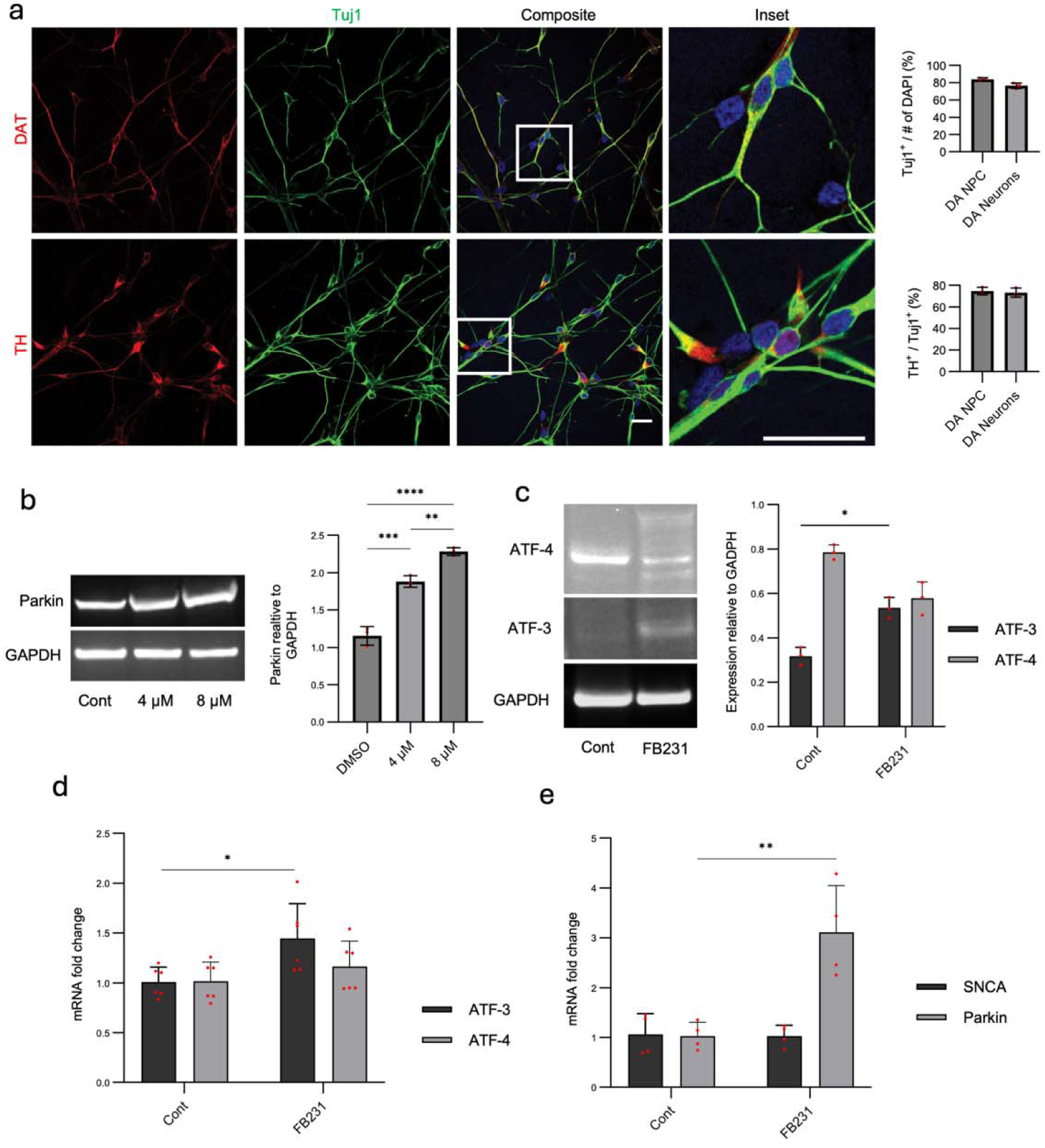
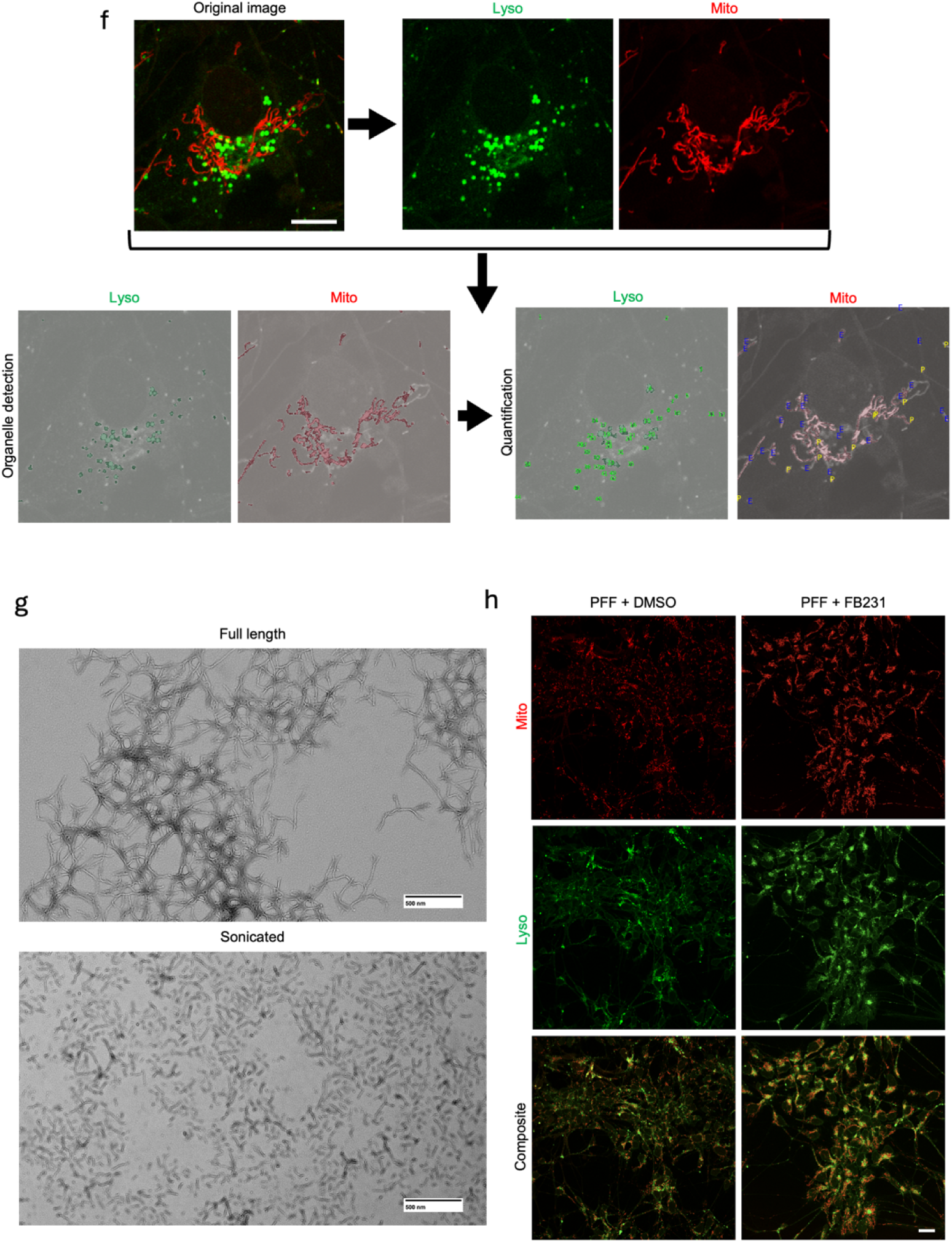
Characterization of iPSC-derived dopaminergic neurons, FB231-upregulation of mitophagy, and PFFs. (**a**) TH and DAT staining in matured iPSC-derived dopaminergic neurons used to assess the efficiency of dopaminergic neuron generation. Quantification of Tuj1-positive neurons divided by the total number of cells (DAPI: 4,6-diamidino-2-phenylindole) yielded the percentage of neuronal cells per imaging field. Scale bar = 20 µm. Additionally, the quantification of TH-positive neurons divided by the total number of neurons (number of Tuj1-positive) yielded the percentage of dopaminergic neurons per imaging field. (**b**) Parkin upregulation in healthy iPSC-derived dopaminergic neurons following 48 h of incubation with 4 µM and 8 µM FB231 or DMSO (Cont). GAPDH was used as a loading control. (**c**) FB231 (4 µM) and DMSO-treated dopaminergic neurons were harvested 36 hours after treatment for Western blot and other experiments. (**d** and **e**) qPCR analysis of the samples used in **c**. (**f**) Analysis pipeline used for detecting and quantifying organelle morphology. Scale bar = 10 µm. (**g**) Transmission electron microscopy (TEM) characterization of recombinant PFFs before and after sonication, via fibril length. (**h**) Large-field Mitotracker (Mito) and Lysotracker (Lyso) staining and colocalization in neurons previously exposed to PFFs at 1 µg/mL for 24 h, then exposed to 4µM FB231 or DMSO for 24 h. Scale bar = 20 µm.

**Supplementary Figure 7.**
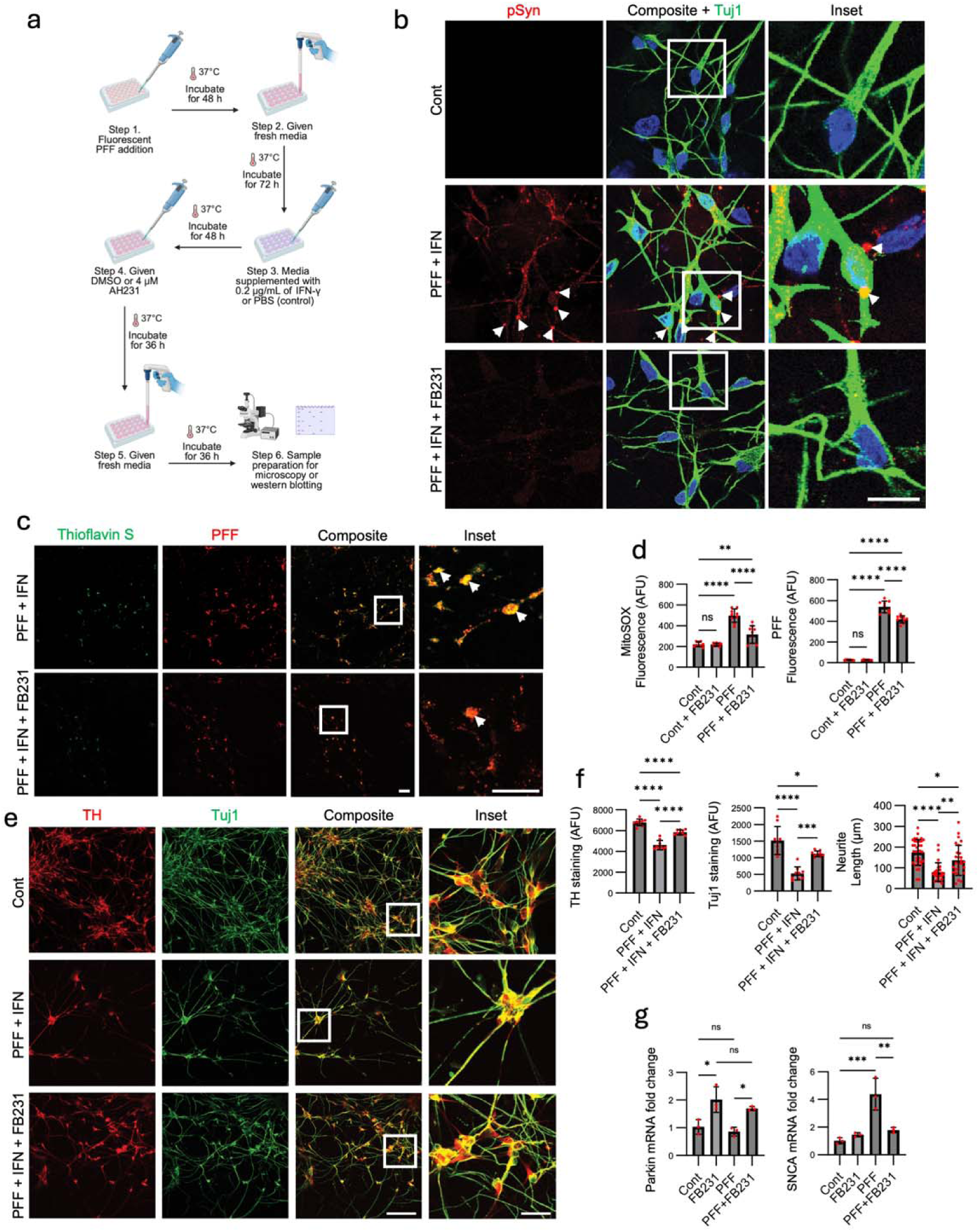
The neuroprotective effects of FB231 in neurons exposed to PFF and IFN-γ. (**a**) Outline of the adapted dual hit treatment regime used in this study. (**b**) Dopaminergic neurons underwent the dual hit treatment regimen for 10 days and then were stained for pSyn. PFF + IFN-γ-treated conditions exhibited high levels of pSyn, with several larger punctae indicating αSyn-positive inclusions (arrowhead). However, when IFN-γ exposure was followed by exposure to FB231, pSyn fluorescence was dramatically reduced. Scale bar = 10 µm. (**c**) Thioflavin S staining of PFF-positive inclusions in PFF + IFN-γ and PFF + IFN-γ + FB231 samples. Scale bar = 20 µm. (**d**) Unbiased quantification of PFF and MitoSOX fluorescence, along with TH and Tuj1 fluorescence, was done for additional experimental and control groups, including control samples treated with FB231, along with PFF-only samples with/without FB231 treatment. (**e**) Neuronal survival and neurite morphology assessed by TH and Tuj1 immunostaining. Scale bar = 100 µm and 25 µm for insets. (**f**) Unbiased quantification of TH and Tuj1 fluorescence and neurite length measured by NeuronJ plugin. (**g**) qPCR analysis of the samples visualized in **e**. All plots show individual data points, the mean, and SD. One-way ANOVA and *post-hoc* Tukey’s test were used for statistical analysis. *p<0.05, **p<0.01, ***p<0.001, ****p<0.001; “ns”, not statistically significant.

## Reference

1 Durcan, T. M. & Fon, E. A. USP8 and PARK2/parkin-mediated mitophagy. Autophagy 11, 428–429 (2015). 10.1080/15548627.2015.1009794

2 Gong, Y. et al. Pan-cancer genetic analysis identifies PARK2 as a master regulator of G1/S cyclins. Nat Genet 46, 588–594 (2014). 10.1038/ng.2981

3 Poole, A. C. et al. The PINK1/Parkin pathway regulates mitochondrial morphology. Proc Natl Acad Sci U S A 105, 1638–1643 (2008). 10.1073/pnas.0709336105

4 Li, H. et al. Longitudinal tracking of neuronal mitochondria delineates PINK1/Parkin-dependent mechanisms of mitochondrial recycling and degradation. Sci Adv 7 (2021). 10.1126/sciadv.abf6580

5 Kitada, T. et al. Mutations in the parkin gene cause autosomal recessive juvenile parkinsonism. Nature 392, 605–608 (1998). 10.1038/33416

6 Marder, K. S. et al. Predictors of parkin mutations in early-onset Parkinson disease: the consortium on risk for early-onset Parkinson disease study. Arch Neurol 67, 731–738 (2010). 10.1001/archneurol.2010.95

7 Nichols, W. C. et al. Linkage stratification and mutation analysis at the Parkin locus identifies mutation positive Parkinson’s disease families. J Med Genet 39, 489–492 (2002). 10.1136/jmg.39.7.489

8 Gong, Y. et al. Pan-Cancer Analysis Links PARK2 to BCL-XL-Dependent Control of Apoptosis. Neoplasia 19, 75–83 (2017). 10.1016/j.neo.2016.12.006

9 Lee, S. B. et al. Parkin Regulates Mitosis and Genomic Stability through Cdc20/Cdh1. Mol Cell 60, 21–34 (2015). 10.1016/j.molcel.2015.08.011

10 Kamienieva, I., Duszynski, J. & Szczepanowska, J. Multitasking guardian of mitochondrial quality: Parkin function and Parkinson’s disease. Transl Neurodegener 10, 5 (2021). 10.1186/s40035-020-00229-8

11 Khan, N. L. et al. Parkin disease: a phenotypic study of a large case series. Brain 126, 1279–1292 (2003). 10.1093/brain/awg142

12 Rub, C., Wilkening, A. & Voos, W. Mitochondrial quality control by the Pink1/Parkin system. Cell Tissue Res 367, 111–123 (2017). 10.1007/s00441-016-2485-8

13 Turrens, J. F. Mitochondrial formation of reactive oxygen species. J Physiol 552, 335–344 (2003). 10.1113/jphysiol.2003.049478

14 Lucking, C. B. et al. Association between early-onset Parkinson’s disease and mutations in the parkin gene. N Engl J Med 342, 1560–1567 (2000). 10.1056/NEJM200005253422103

15 Dawson, T. M. & Dawson, V. L. The role of parkin in familial and sporadic Parkinson’s disease. Mov Disord 25, S32–39 (2010). 10.1002/mds.22798

16 Jiang, Y. et al. Parkin is the most common causative gene in a cohort of mainland Chinese patients with sporadic early-onset Parkinson’s disease. Brain Behav 10, e01765 (2020). 10.1002/brb3.1765

17 Wang, C. et al. Stress-induced alterations in parkin solubility promote parkin aggregation and compromise parkin’s protective function. Hum Mol Genet 14, 3885–3897 (2005). 10.1093/hmg/ddi413

18 Wong, E. S. et al. Relative sensitivity of parkin and other cysteine-containing enzymes to stress-induced solubility alterations. J Biol Chem 282, 12310–12318 (2007). 10.1074/jbc.M609466200

19 Sunico, C. R. et al. S-Nitrosylation of parkin as a novel regulator of p53-mediated neuronal cell death in sporadic Parkinson’s disease. Mol Neurodegener 8, 29 (2013). 10.1186/1750-1326-8-29

20 Lonskaya, I., Hebron, M. L., Algarzae, N. K., Desforges, N. & Moussa, C. E. Decreased parkin solubility is associated with impairment of autophagy in the nigrostriatum of sporadic Parkinson’s disease. Neuroscience 232, 90–105 (2013). 10.1016/j.neuroscience.2012.12.018

21 Shlevkov, E., Kramer, T., Schapansky, J., LaVoie, M. J. & Schwarz, T. L. Miro phosphorylation sites regulate Parkin recruitment and mitochondrial motility. Proc Natl Acad Sci U S A 113, E6097–E6106 (2016). 10.1073/pnas.1612283113

22 Shlevkov, E. et al. Discovery of small-molecule positive allosteric modulators of Parkin E3 ligase. iScience 25, 103650 (2022). 10.1016/j.isci.2021.103650

23 Sauve, V. et al. Activation of parkin by a molecular glue. Nat Commun 15, 7707 (2024). 10.1038/s41467-024-51889-3

24 Rosencrans, W. M. et al. Putative PINK1/Parkin activators lower the threshold for mitophagy by sensitizing cells to mitochondrial stress. Sci Adv 11, eady0240 (2025). 10.1126/sciadv.ady0240

25 Tompkins, M. M. & Hill, W. D. Contribution of somal Lewy bodies to neuronal death. Brain Research 775, 24–29 (1997). 10.1016/S0006-8993(97)00874-3

26 Vives-Bauza, C. et al. PINK1-dependent recruitment of Parkin to mitochondria in mitophagy. Proc Natl Acad Sci U S A 107, 378–383 (2010). 10.1073/pnas.0911187107

27 Norris, K. L. et al. Convergence of Parkin, PINK1, and alpha-Synuclein on Stress-induced Mitochondrial Morphological Remodeling. J Biol Chem 290, 13862–13874 (2015). 10.1074/jbc.M114.634063

28 Wu, W. et al. PINK1-Parkin-Mediated Mitophagy Protects Mitochondrial Integrity and Prevents Metabolic Stress-Induced Endothelial Injury. PLoS One 10, e0132499 (2015). 10.1371/journal.pone.0132499

29 Ishikawa, E. T. & Cancelas, J. A. Lack of communication rusts and ages stem cells. Cell Cycle 11, 3149–3150 (2012). 10.4161/cc.21589

30 Taniguchi Ishikawa, E., et al. Connexin-43 prevents hematopoietic stem cell senescence through transfer of reactive oxygen species to bone marrow stromal cells. Proc Natl Acad Sci U S A 109, 9071–9076 (2012). 10.1073/pnas.1120358109

31 Tseng, P. Y. & Hoon, M. A. Oncostatin M can sensitize sensory neurons in inflammatory pruritus. Sci Transl Med 13, eabe3037 (2021). 10.1126/scitranslmed.abe3037

32 Sharanek, A. et al. OSMR controls glioma stem cell respiration and confers resistance of glioblastoma to ionizing radiation. Nat Commun 11, 4116 (2020). 10.1038/s41467-020-17885-z

33 Chang, S. H., Hwang, C. S., Yin, J. H., Chen, S. D. & Yang, D. I. Oncostatin M-dependent Mcl-1 induction mediated by JAK1/2-STAT1/3 and CREB contributes to bioenergetic improvements and protective effects against mitochondrial dysfunction in cortical neurons. Biochim Biophys Acta 1853, 2306–2325 (2015). 10.1016/j.bbamcr.2015.05.014

34 Badia-i-Mompel, P. et al. decoupleR: ensemble of computational methods to infer biological activities from omics data. Bioinformatics Advances 2, vbac016 (2022). 10.1093/bioadv/vbac016

35 Seale, P. et al. Transcriptional control of brown fat determination by PRDM16. Cell Metab 6, 38–54 (2007). 10.1016/j.cmet.2007.06.001

36 He, L. et al. PRDM16 regulates a temporal transcriptional program to promote progression of cortical neural progenitors. Development 148 (2021). 10.1242/dev.194670

37 Friedrich, T., Tavraz, N. N. & Junghans, C. ATP1A2 Mutations in Migraine: Seeing through the Facets of an Ion Pump onto the Neurobiology of Disease. Front Physiol 7, 239 (2016). 10.3389/fphys.2016.00239

38 Malloy, C., Ahern, M., Lin, L. & Hoffman, D. A. Neuronal Roles of the Multifunctional Protein Dipeptidyl Peptidase-like 6 (DPP6). Int J Mol Sci 23 (2022). 10.3390/ijms23169184

39 Quintero-Rivera, F., Chan, A., Donovan, D. J., Gusella, J. F. & Ligon, A. H. Disruption of a synaptotagmin (SYT14) associated with neurodevelopmental abnormalities. Am J Med Genet A **143A**, 558–563 (2007). 10.1002/ajmg.a.31618

40 Shimura, H. et al. Ubiquitination of a New Form of α-Synuclein by Parkin from Human Brain: Implications for Parkinson’s Disease. Science 293, 263–269 (2001). 10.1126/science.1060627

41 Kim, S. et al. Transneuronal Propagation of Pathologic alpha-Synuclein from the Gut to the Brain Models Parkinson’s Disease. Neuron 103, 627–641 e627 (2019). 10.1016/j.neuron.2019.05.035

42 Van Den Berge, N. & Ulusoy, A. Animal models of brain-first and body-first Parkinson’s disease. Neurobiol Dis 163, 105599 (2022). 10.1016/j.nbd.2021.105599

43 Challis, C. et al. Gut-seeded alpha-synuclein fibrils promote gut dysfunction and brain pathology specifically in aged mice. Nat Neurosci 23, 327–336 (2020). 10.1038/s41593-020-0589-7

44 Scholz, J., Niibori, Y., P, W. F. & J, P. L. Rotarod training in mice is associated with changes in brain structure observable with multimodal MRI. Neuroimage 107, 182–189 (2015). 10.1016/j.neuroimage.2014.12.003

45 Shiotsuki, H. et al. A rotarod test for evaluation of motor skill learning. J Neurosci Methods 189, 180–185 (2010). 10.1016/j.jneumeth.2010.03.026

46 Ogawa, N., Hirose, Y., Ohara, S., Ono, T. & Watanabe, Y. A simple quantitative bradykinesia test in MPTP-treated mice. Res Commun Chem Pathol Pharmacol 50, 435–441 (1985).

47 Jeong, D. et al. Modulation of gut microbiota and increase in fecal water content in mice induced by administration of Lactobacillus kefiranofaciens DN1. Food Funct 8, 680–686 (2017). 10.1039/c6fo01559j

48 Anderson, J. P. et al. Phosphorylation of Ser-129 is the dominant pathological modification of alpha-synuclein in familial and sporadic Lewy body disease. J Biol Chem 281, 29739–29752 (2006). 10.1074/jbc.M600933200

49 Dent, S. E. et al. Phosphorylation of the aggregate-forming protein alpha-synuclein on serine-129 inhibits its DNA-bending properties. J Biol Chem 298, 101552 (2022). 10.1016/j.jbc.2021.101552

50 Bayati, A. et al. Modeling Parkinson’s disease pathology in human dopaminergic neurons by sequential exposure to alpha-synuclein fibrils and proinflammatory cytokines. Nat Neurosci 27, 2401–2416 (2024). 10.1038/s41593-024-01775-4

51 Koncha, R. R., Ramachandran, G., Sepuri, N. B. V. & Ramaiah, K. V. A. CCCP-induced mitochondrial dysfunction - characterization and analysis of integrated stress response to cellular signaling and homeostasis. FEBS J 288, 5737–5754 (2021). 10.1111/febs.15868

52 Giedt, R. J. et al. Computational imaging reveals mitochondrial morphology as a biomarker of cancer phenotype and drug response. Sci Rep 6, 32985 (2016). 10.1038/srep32985

53 Murillo-Gonzalez, F. E., Garcia-Aguilar, R., Vega, L. & Elizondo, G. Regulation of Parkin expression as the key balance between neural survival and cancer cell death. Biochem Pharmacol 190, 114650 (2021). 10.1016/j.bcp.2021.114650

54 Han, R. et al. Deficiency of parkin causes neurodegeneration and accumulation of pathological alpha-synuclein in monkey models. J Clin Invest 134 (2024). 10.1172/JCI179633

55 Cooper, J. F. et al. Activation of the mitochondrial unfolded protein response promotes longevity and dopamine neuron survival in Parkinson’s disease models. Sci Rep 7, 16441 (2017). 10.1038/s41598-017-16637-2

56 Chung, K. K. et al. Parkin ubiquitinates the alpha-synuclein-interacting protein, synphilin-1: implications for Lewy-body formation in Parkinson disease. Nat Med 7, 1144–1150 (2001). 10.1038/nm1001-1144

57 Khandelwal, P. J. et al. Parkinson-related parkin reduces α-Synuclein phosphorylation in a gene transfer model. Molecular Neurodegeneration 5, 47 (2010). 10.1186/1750-1326-5-47

58 Tufail, M., Jiang, C. H. & Li, N. Altered metabolism in cancer: insights into energy pathways and therapeutic targets. Mol Cancer 23, 203 (2024). 10.1186/s12943-024-02119-3

59 Ni, A. & Ernst, C. Evidence That Substantia Nigra Pars Compacta Dopaminergic Neurons Are Selectively Vulnerable to Oxidative Stress Because They Are Highly Metabolically Active. Front Cell Neurosci 16, 826193 (2022). 10.3389/fncel.2022.826193

60 Danussi, C. et al. Atrx inactivation drives disease-defining phenotypes in glioma cells of origin through global epigenomic remodeling. Nature Communications 9, 1057 (2018). 10.1038/s41467-018-03476-6

61 Joung, J. et al. Genome-scale CRISPR-Cas9 knockout and transcriptional activation screening. Nature Protocols 12, 828–863 (2017). 10.1038/nprot.2017.016

62 Cai, W. et al. Melanocortin 1 receptor activation protects against alpha-synuclein pathologies in models of Parkinson’s disease. Mol Neurodegener 17, 16 (2022). 10.1186/s13024-022-00520-4

63 Cai, W., Feng, D., Schwarzschild, M. A., McLean, P. J. & Chen, X. Bimolecular Fluorescence Complementation of Alpha-synuclein Demonstrates its Oligomerization with Dopaminergic Phenotype in Mice. EBioMedicine 29, 13–22 (2018). 10.1016/j.ebiom.2018.01.035

64 Granucci, E. J. et al. Cromolyn sodium delays disease onset and is neuroprotective in the SOD1(G93A) Mouse Model of amyotrophic lateral sclerosis. Sci Rep 9, 17728 (2019). 10.1038/s41598-019-53982-w

65 Xu, K. et al. Estrogen Prevents Neuroprotection by Caffeine in the Mouse 1-Methyl-4-Phenyl-1,2,3,6-Tetrahydropyridine Model of Parkinson’s Disease. The Journal of Neuroscience 26, 535 (2006). 10.1523/JNEUROSCI.3008-05.2006

66 Xu, K. et al. Neuroprotection by caffeine in the MPTP model of parkinson’s disease and its dependence on adenosine A2A receptors. Neuroscience 322, 129–137 (2016). 10.1016/j.neuroscience.2016.02.035

67 Cipriani, S., Bakshi, R. & Schwarzschild, M. A. Protection by inosine in a cellular model of Parkinson’s disease. Neuroscience 274, 242–249 (2014). 10.1016/j.neuroscience.2014.05.038

68 Meijering, E. et al. Design and validation of a tool for neurite tracing and analysis in fluorescence microscopy images. Cytometry A 58, 167–176 (2004). 10.1002/cyto.a.20022

69 Volpicelli-Daley, L. A., Luk, K. C. & Lee, V. M. Addition of exogenous alpha-synuclein preformed fibrils to primary neuronal cultures to seed recruitment of endogenous alpha-synuclein to Lewy body and Lewy neurite-like aggregates. Nat Protoc 9, 2135–2146 (2014). 10.1038/nprot.2014.143

70 Bayati, A. et al. Rapid macropinocytic transfer of alpha-synuclein to lysosomes. Cell Rep 40, 111102 (2022). 10.1016/j.celrep.2022.111102

71 Bayati, A. et al. Visualization of alpha-synuclein trafficking via nanogold labeling and electron microscopy. STAR Protoc 4, 102113 (2023). 10.1016/j.xpro.2023.102113

72 Maneca, D.-L. et al. Production of Recombinant α Synuclein Monomers and Preformed Fibrils (PFFs). Zenodo (2019). 10.5281/zenodo.3738335

73 Srivastava, P. et al. Peripheral MC1R Activation Modulates Immune Responses and is Neuroprotective in a Mouse Model of Parkinson’s Disease. J Neuroimmune Pharmacol 18, 704–717 (2023). 10.1007/s11481-023-10094-7

74 Noorian, A. R. et al. Alpha-synuclein transgenic mice display age-related slowing of gastrointestinal motility associated with transgene expression in the vagal system. Neurobiol Dis 48, 9–19 (2012). 10.1016/j.nbd.2012.06.005

